# Magnesium maintains length of circadian period in *Arabidopsis thaliana*

**DOI:** 10.1101/2020.05.14.096537

**Authors:** J. Romário F. de Melo, Annelie Gutsch, Joëlle De Caluwé, Jean-Christophe Leloup, Didier Gonze, Christian Hermans, Alex A.R. Webb, Nathalie Verbruggen

## Abstract

The circadian clock coordinates the physiological response of a biological system to day and night rhythms through complex loops of transcriptional/ translational regulation. It can respond to external stimuli and adjust generated circadian oscillations accordingly to keep an endogenous period close to 24 h. To date, the interaction between nutritional status and circadian rhythms in plants is poorly understood. Magnesium (Mg) is essential for numerous biological processes in plants and its homeostasis is crucial to maintain optimal development and growth. Magnesium deficiency in young *Arabidopsis thaliana* seedlings increased the circadian period of *pCCA1:LUC* oscillations and dampened its amplitude in constant light in a dose-dependent manner. Although circadian period increase by Mg deficiency was light dependent, it did not depend on active photosynthesis. Mathematical modelling of the Mg input to the circadian clock reproduced the experimental increase of the circadian period and suggested that Mg is likely to affect global transcription/translation levels rather than a single component of the circadian oscillator. The model prediction was supported by a synergistic interaction between Mg deficiency and cyclohexamide, an inhibitor of translation. These findings suggest that proper Mg supply is required to support proper timekeeping in plants.

**One sentence summary:** Magnesium maintains the circadian period in Arabidopsis seedlings and interferes with the circadian oscillator most likely through translational mechanisms.

## Introduction

Magnesium (Mg) is one of the most abundant elements in the Earth’s crust (Clark and Washington, 1924; Fleischer, 1954) and in sea water (Culkin and Cox, 1966). It plays many roles in the metabolism of living organisms, such as maintaining ribosome structure (Akanuma et al., 2018), being necessary for the active form of ATP (Fish et al., 1983; Wang et al., 1995) as well as being a co-factor and allosteric modulator for numerous enzymes (Cowan, 1998). In plants, Mg is vital to the photosynthetic machinery (Levitt, 1954) and CO_2_ assimilation (Hauer-Jákli and Tränkner, 2019). Therefore, imbalances in plant Mg status are likely to cause disorders from cellular to organismal levels. Despite its importance in cellular and organismal metabolic processes across all kingdoms, Mg still does not garner as much attention as other nutrients, such as Ca, N, Zn or Fe (Hermans et al., 2013).

Plants require usually between 1.5 – 3.5 mg g^−1^ dry weight for optimal growth (Grzebisz et al., 2009; Römheld et al., 2012). A Mg supply below 1 – 2 mg g-1 leaf dry weight marks the onset of Mg deficiency (Hermans et al. 2004, Hermans and Verbruggen et al., 2005, Ding et al., 2006). Impaired partitioning of soluble sugars leading to starch accumulation in source leaves are the first sign of Mg deficiency before defects of photosynthetic activity occur (Cakmak et al., 1994; Hermans et al., 2005; Hermans and Verbruggen, 2005). Typical long-term symptoms of Mg starvation in higher plants are interveinal leaf chlorosis, limited growth and altered biomass allocation between plant organs (Verbruggen and Hermans, 2013; Hauer-Jákli and Tränkner, 2019). Transcriptomic studies in *Arabidopsis. thaliana* identified *CATION EXCHANGER 3* (*CAX3*) as a suitable molecular marker to monitor the Mg status because it responds to Mg availability before the first visible signs of deficiency or excess occur (Hermans et al., 2010a; Kamiya et al., 2012). Transcriptomic studies further revealed that both early, as well as long-term Mg deficiency, altered the expression of genes involved in processes regulated by the circadian oscillator (Hermans et al., 2010a; Hermans et al., 2010b)

Daily biological rhythms in plants are regulated by the circadian clock, which runs in a close-to 24-hour cycle synchronized to environmental cues such as light and temperature. The circadian clock is maintained by endogenous rhythms of gene expression regulated by transcriptional-translational feedback loops that influence growth, development, flowering time, responses to biotic and abiotic stresses to promote plant fitness (Green et al., 2002; Harmer, 2009; Greenham and McClung, 2015). The core components of the central oscillator in plants are the dawn-phased genes *CIRCADIAN CLOCK-ASSOCIATED 1* (*CCA1*) and *LATE ELONGATED HYPOCOTYL* (*LHY*), the morning genes *PSEUDO-RESPONSE REGULATOR 9 (PRR9), PRR7* and *PRR5*, the dusk-phased gene *TIMING OF CAB EXPRESSION 1* (*TOC1*) and the evening-complex composed of *EARLY FLOWERING 3 (ELF3), ELF4*, and *LUX ARRYHTHMO* (*LUX*) (McClung, 2006; Webb et al., 2019). Altered expression of one or more of these oscillator components changes the circadian period, amplitude, phase and can lead to complete arrhythmia of the endogenous oscillator, which affects plant growth and development (Hicks et al., 1996; Dunlap, 1999; Alabadí et al., 2002; Webb, 2003). Therefore, optimal functioning of the circadian system relies on sensing and integrating internal as well as external signals in order to maintain an endogenous timekeeping mechanism capable of accurately anticipating environmental fluctuations (Dodd et al., 2005; Hotta et al., 2007; Robertson et al., 2009; Hsu and Harmer, 2014). Such external signals can be the nutritional status that cross talks with the circadian clock (Hong et al., 2013) and imbalanced nutritional homeostasis can interfere with circadian timekeeping. Experiments, which establish the effect of stimuli to change the phase of the circadian oscillator at different times of day, so called phase response curve (PRC) experiments, using pulses of nitrogen (N) support the feedback of N status to the circadian oscillator (Gutiérrez et al., 2008). Other studies demonstrated that N deficiency shortened-circadian period in the photosynthetic dinoflagellate *Gonyaulax polyedra* (Sweeney and Folli, 1984; Haydon et al., 2015). Excess of copper affects amplitude and phase of *CCA1* and *LHY* expression (Andrés-Colás et al., 2010) while iron deficiency lengthens the circadian period (Chen et al., 2013; Salomé et al., 2013). Furthermore, early or long-term Mg deficiency alters the expression of Arabidopsis circadian oscillator genes suggesting a link between Mg homeostasis and the circadian clock (Hermans et al., 2010a; Hermans et al., 2010b). While no detailed study describes the interplay between Mg and circadian rhythms in plants, recent findings suggest a key role for Mg in the timekeeping system in *Ostreococcus tauri* (Feeney et al., 2016), a single-celled alga that shares a common ancestor with higher plants.

We show that Mg deficiency dose-dependently lengthens the circadian period of core oscillator genes in *A. thaliana*, which is independent from fully functional photosynthesis. A comparable period lengthening was reproduced with mathematical modelling when a global impact of Mg on transcription and translation was simulated. Our findings demonstrate that endogenous rhythms in plants strongly depend on nutritional status.

## Results

### External Mg concentrations dose-dependently increases period under continuous light and impact circadian phase in light/dark cycles

Luciferase (LUC)-based reporters were used to examine circadian rhythms in *A. thaliana* seedlings that were germinated and entrained under different Mg concentrations ranging from 5 μM to 1,500 μM and thereafter released into continuous light (LL). Magnesium depletion lengthened the circadian period of *pCCA1:LUC* activity by almost 5 h on average when compared with Mg-replete controls: τ = 28.77 h *vs*. 24.34 h, respectively, in a dose-dependent manner (r = −0.99, p < 0.001; Fig 1*A*-*B*). Increasing period was associated with reduced amplitude of *pCCA1:LUC* oscillations and an increased relative amplitude error (RAE) (Fig 1*A*, *C*). Results were confirmed also with the reporter lines *pPRR7:LUC* and *pTOC:LUC*, whereby the effect of Mg deficiency was more pronounced in the presence of 1 % (w/v) sucrose for all reporter lines tested (Fig. S1). Magnesium deficiency led to reduced growth; the fresh biomass of 15 days old seedlings was about 23 % less than the biomass of seedlings grown under sufficient Mg supply (Fig 1*D-E*). Dry biomass was significantly reduced (*P* < 0.001) and the internal Mg status of seedlings was significantly lower (*P* < 0.001) when external Mg was limited (Fig 1*F-G*).

**Fig. 1.**
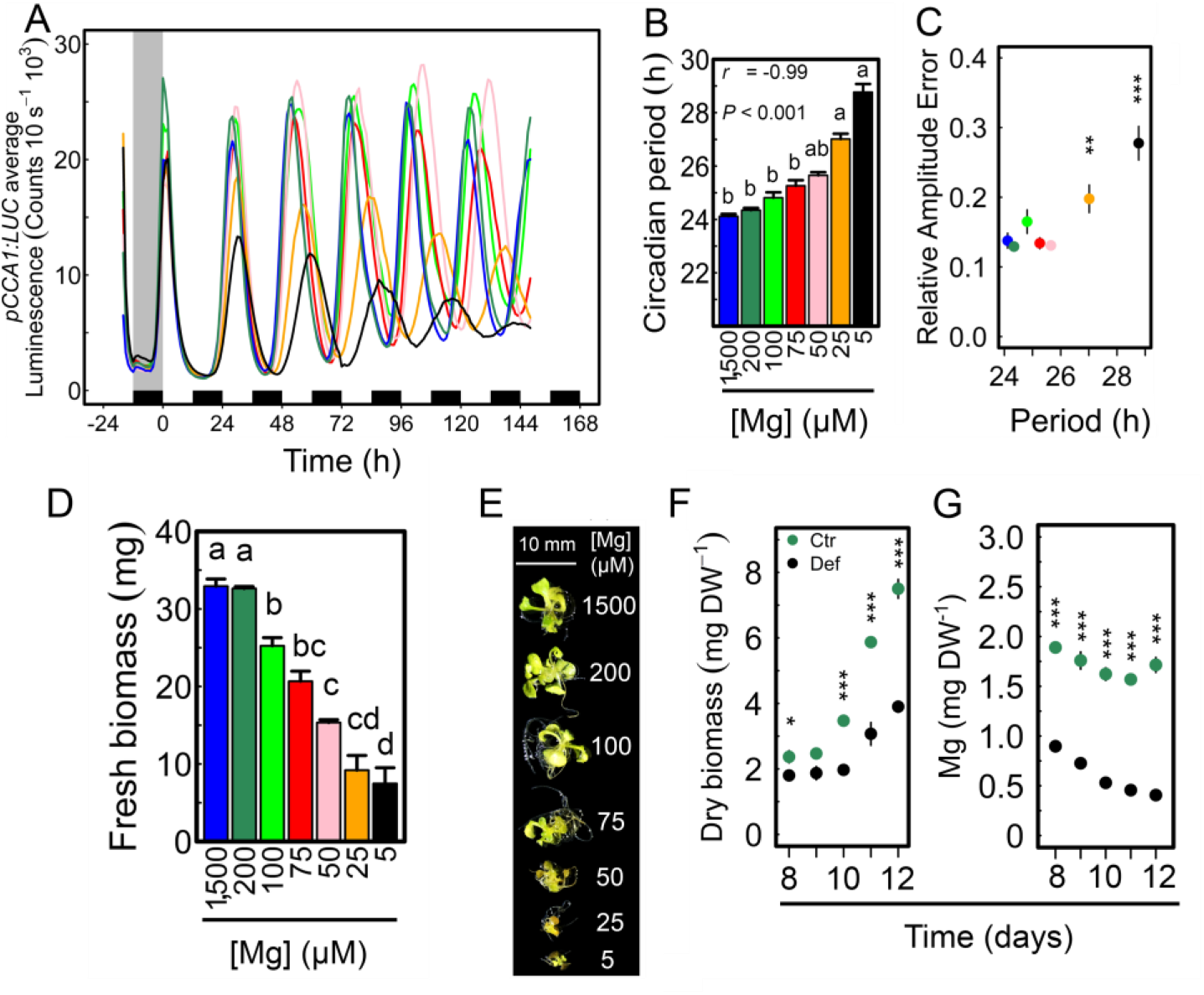
Limiting external Mg availability alters circadian oscillations of *CCA1:LUC*. (*A*) Mean circadian oscillations of *pCCA1:LUC* in LL conditions after being entrained to 12/12-h light/dark cycles for eight days on media supplied with different Mg concentrations. (*B*) *pCCA1:LUC* period estimates in h under LL. (*C*) relative amplitude error of oscillations (mean ± SEM, *n* = 12) of *pCCA1:LUC*. (*D*) Fresh biomass and (*E*) morphological phenotype of seedlings at the end of the experiment. (*F*) dry biomass and (*G*) Mg concentration in plant tissue [mean ± SEM, *n* = 3 (1 = 15 pooled seedlings)] of 12-day-old seedlings cultivated in light/dark cycles on control (Ctr.: 200 μM Mg) and deficient media (Def.: 5 μM Mg). Significance was verified by (*B*) Spearman’s rho correlation coefficient and Kruskal-Wallis Rank Sum Test followed by Nemenyi post-hoc test, (*C*) One-Way ANOVA followed by Tukey HSD post-hoc test and (*F*-*G*) Two-Sample Student’s *t*-test 95% C.I. (different letters indicate significance at the level of *P* < 0.05. Asterisks represent significance at * *P* < 0.05, ** *P* < 0.01, and *** *P* < 0.001).

To test if increased circadian period is linked to growth inhibition provoked by Mg shortage, seedlings were grown under N deficiency, which is also a major macronutrient. Seedlings fully supplied (10 mM) or starved (0.01 mM) with N had a similar free-running circadian period under either conditions: τ = 24.3 h; *vs*. 24.3 h (Fig. S2*A*-*B*). Yet, RAE of *pCCA1* oscillations was affected by low-N supply (Fig. S2*C-D*) and seedlings had morphology and size characteristic of severe N deficiency (Figure S2*E*). These results show that severe growth inhibition induced by nutrient deficiency and circadian period lengthening are not correlated.

The effect of Mg nutrition on circadian phase was determined in light/dark cycles (12/12) in seedlings sown on different concentrations of Mg (Fig. 2). In light/dark cycles *TOC1* is often timed to dusk, and consistent with that no effect of Mg or sucrose on the phase of *pTOC1:LUC* activity was found (Fig. 2*A*). The phase of peak activity of both *pCCA1:LUC* and *pPRR7:LUC* was advanced by more than one hour by exogenous sucrose similar to previous observations (Fig. 2*A*) (Haydon et al., 2013; Frank et al., 2018). In the absence of added sucrose depletion of Mg resulted in a phase delay of both *pCCA1:LUC* and *pPRR7:LUC* activity, however, in the presence of added 1% sucrose only the phase of *pPRR7:LUC* activity was sensitive to Mg status (Fig. 2*A*). It appears that *PRR7* is the most plastic component during the circadian cycle of Arabidopsis responding to Mg depletion. To examine this observation further, *pPRR7:LUC* activity was measured in a 12/12 hour light/dark cycle over the course of four days under different Mg supply. A dose dependent effect of Mg on the phase of *pPPR7:LUC* was strongest in the presence of 1% added sucrose but was also observed in the absence of added sucrose (Fig. 2*B-C*). We conclude that there is a plastic response of the circadian oscillator to Mg in diel cycles of light and dark, with *PRR7* being the most plastic component having a phase delay in response to low Mg. Despite the phase delay of *pPRR7:LUC* due to Mg shortage, the phase of the circadian oscillator was not altered and the estimated period was approximately 24 hours under all concentrations tested due to the forcing light/dark cycle (Fig. S3).

**Fig. 2.**
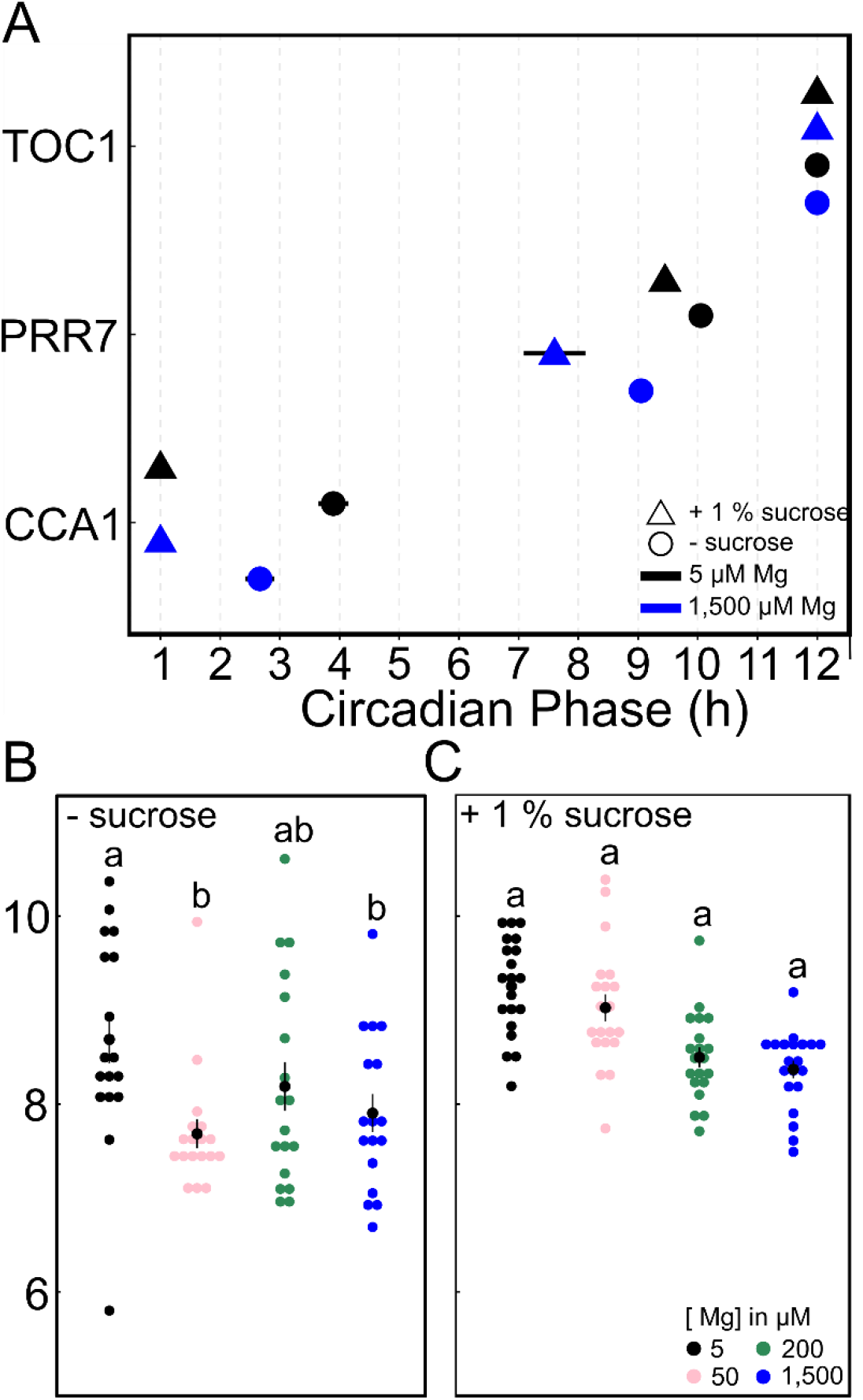
Mg deficiency delays the circadian phase of *PRR7*. (*A*) Circadian phase of peak expression (mean ± SEM) in 12/12 hour light/dark cycle of *pCCA1:LUC, pPRR7:LUC* and *pTOC1:LUC* in 11-day-old Col-0 WT Arabidopsis seedlings (n=20) in the absence or presence of 1 % sucrose. Circadian phase of *pPRR7:LUC* peak expression in a four day 12/12 hour light/dark cycle under different Mg supply in the (*B*) absence or (*C*) presence of 1 % sucrose. Significance was determined by a Wilcoxon Rang Sum test. Different letters indicate significance at the level of *P* ≤ 0.05.

### Induction of a Mg deficiency-marker is congruent with circadian clock alteration

Neither a difference in circadian oscillations nor an effect on plant growth was observed when seedlings were supplied with 200 μM Mg or 1,500 μM Mg (Fig. 1*A-E*). Therefore, it was tested whether a more rapid Mg-deficient status in plants could be induced upon pre-cultivation on 200 μM Mg in comparison to the usually used 1,500 μM Mg. In fact, metabolically available Mg concentrations in plant cells are relatively high (15-25 mM) and the vacuolar storage is reported to range from 5 to 80 mM (Hermans et al., 2013). Seedlings were entrained on either 1,500 μM or 200 μM Mg and thereafter transferred to Mg-deficient medium (5 μM) for another five days in LL (Fig. S4*A*). Seedlings transferred to 5 μM Mg were pale and produced significantly less biomass only when pre-cultured on 200 μM Mg (Fig. S4*B-C*). *CAX3*-transcript levels were low and comparable between 1,500 μM and 200 μM Mg control conditions (Fig. S4D). In seedlings subjected to 5 μM after being pre-cultured on 1,500 μM Mg, *CAX3* transcript levels increased 72 h after Mg removal (Fig. 3*A*). But, in seedlings pre-cultured on 200 μM Mg *CAX3* expression was already induced after 48 h of treatment (Fig. 3*B*). Apparently, a supply with 1,500 μM Mg provides enough Mg storage to prevent Mg deficiency within the first three days of deprivation. However, when seedlings were entrained on 200 μM Mg the removal of Mg induced a more rapid and severe response towards deficiency stress (Fig. 3*B*, Fig. S4*B-C*). Therefore, 200 μM Mg was chosen as a new control concentration to entrain seedlings and to investigate the time course of clock alteration and induction of Mg deficiency.

**Fig. 3.**
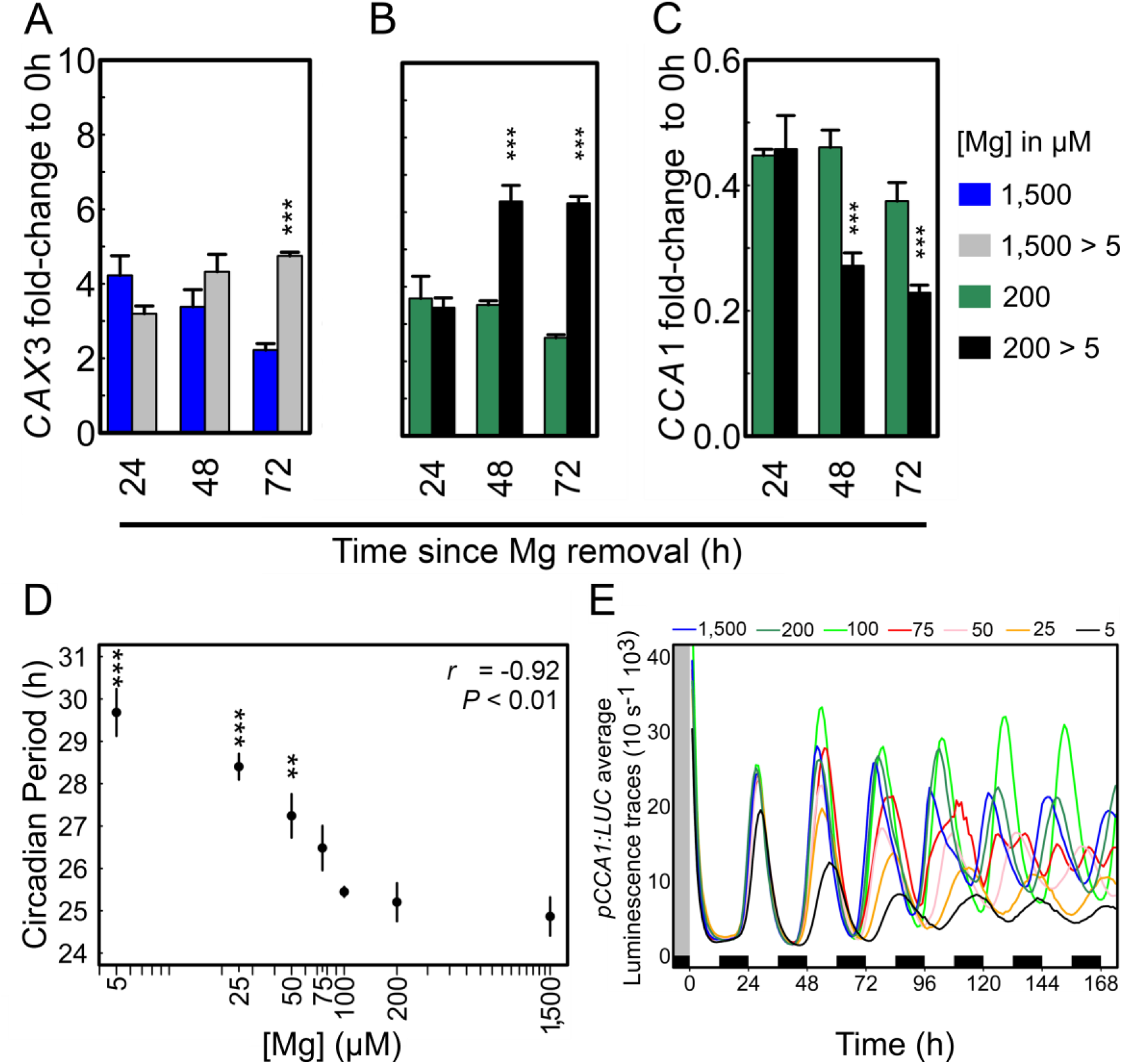
Oscillator alteration occurs concomitantly with expression of the Mg deficiency marker CAX3. Seedlings entrained for eight days to 12/12-h light/dark cycles on medium with 200 or 1,500 μM Mg were transferred to fresh media either deficient in Mg (5 μM) or fully supplied and released in continuous light. (*A*) *CAX3* mRNA expression [mean ± SEM, *n* = 3 (1 = 30 pooled seedlings)] after transfer from 1,500 μM Mg, (B-C) *CAX3* mRNA expression and *CCA1* mRNA expression [mean ± SEM, *n* = 3 (1 = 30 pooled seedlings)] after transfer from 200 μM Mg, (*D*) correlation between estimated circadian period and external Mg concentration, (*E*) average luminescence traces of *pCCA1:LUC* activity. Significance was verified by Pearson’s Product Moment correlation coefficient (*P* < 0.01) and One-Way ANOVA followed by Tukey HSD post-hoc or Two-Sample Student’s *t*-test at a 95% confidence interval (asterisks represent significance at * *P* < 0.05, ** *P* < 0.01, and *** *P* < 0.001).

In seedlings entrained on medium with 200 μM Mg, mean *CCA1* transcript levels decreased 2-fold 48-72 h after being transferred to 5 μM Mg (Fig. 3C), which coincided with the significant increase in *pCCA1:LUC* period induced by Mg deficiency (Fig 3*D*). Severe Mg deficiency (5 μM Mg) caused a circadian response one day in LL after the transfer (Fig. 3*E*). Thereby, *CCA1* expression decrease appeared at the same time point as the expression increase of *CAX3* mRNA, an indicator for Mg status of the plant (Fig. 3*B*).

### Light plays a critical role on the circadian effects of Mg deficiency

Some circadian alterations are manifested in a light conditional manner (Hicks et al., 1996). To test the role of light in the response of the circadian oscillations in *A. thaliana* to Mg, seedlings were entrained to either 8/16-h light/dark cycles (short days: SD) or 16/8-h light/dark cycles (long days: LD). In SD, the biomass of Mg deficient seedlings was 59 % reduced in comparison to the respective control seedlings and in LD the reduction was 81 % (Fig. 4*A*). Circadian oscillations of *pCCA1:LUC* were monitored after release into LL. Similar to 12/12 h light/dark entrainment, Mg deficiency increased the period and RAE of *pCCA1:LUC* oscillations for both entrained photoperiods. However, the effect on period was stronger in seedlings entrained to LD conditions (Fig. 4*B-D*). In fact, Mg deficiency had no effect on period after plants were released to constant darkness (DD) (Def: τ = 24.16 h *vs*. Ctr: τ = 24.17 h) (Fig. 4*E-F*) that suggests a light dependence of the response of the oscillator to Mg deficiency. After seedlings were released into DD circadian activity was dampened after 48 h in darkness (Fig. 4*E*) but rhythms sustained for the duration of the experiment, which probably results from the presence of 1 % sucrose in the culture medium (Table S1) (Dalchau et al., 2011).

**Fig. 4.**
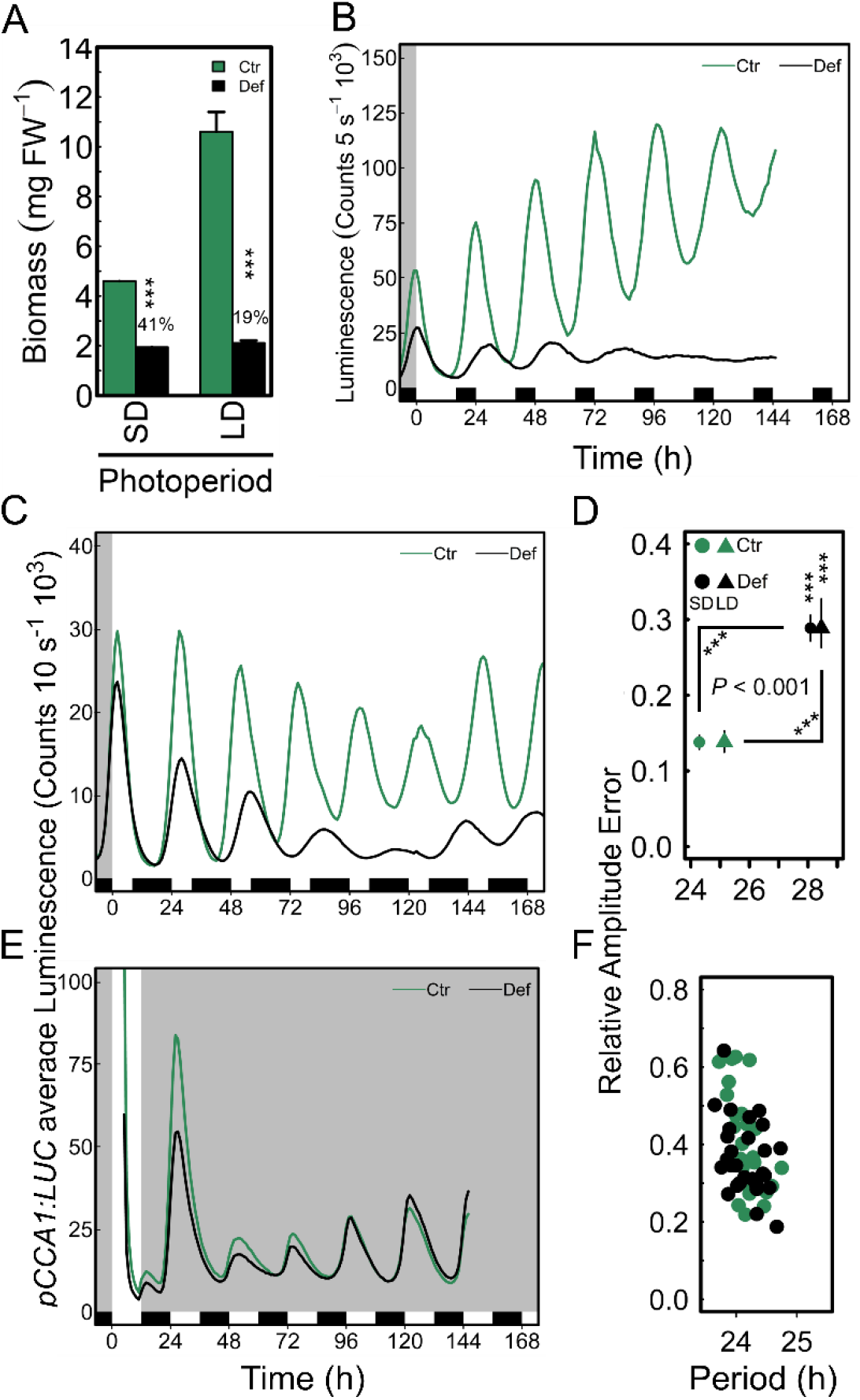
Increase of circadian period due to Mg deficiency is light dependent. (*A*) fresh weight (FW) biomass (mean ± SEM, *n* = 3-4 [1 = 4 pooled seedlings]) of seedlings entrained to long day (LD) 16/8 light/dark cycles and short day (SD) 8/16 h light/dark cycles. *pCCA1:LUC* average luminescence traces of seedlings entrained for eight days to (*B*) LD, (*C*) SD on Mg sufficient (Ctr.: 200 μM) and deficient (Def.: 5 μM) medium before released into LL, (*D*) relative amplitude error of LD and SD (mean ± SEM, *n* = 12-48). Diagonal-oriented asterisks represent significance for the circadian period, vertical-oriented asterisks represent significance for relative amplitude error. (*E*) *pCCA1:LUC* average luminescence traces of seedlings entrained for eight days to a 12/12 h light/dark cycles on Ctr. and Def. media before released into DD, (*F*) respective relative amplitude error. Significance was verified by Two-Sample Student’s *t*-test at 95% confidence interval (*** *P* < 0.001).

### Photosynthesis inhibition does not explain the response of the circadian oscillator to Mg deficiency

To investigate whether the effect of Mg depletion was due to an inhibition of photosynthesis, we examined the effects of Mg in the presence or absence of 3-(3,4-dichlorophenyl)-1, 1-dimethylurea (DCMU), an inhibitor of the photochemical activity of photosystem II. Experiments were performed in the presence of 1 % sucrose as we observed that Mg depletion in the presence of sucrose has a strong effect on circadian period. Therefore, including sucrose in the medium allows to examine direct effects of photosynthetic inhibition, such as retrograde signalling, rather than effects caused by sugar depletion due to inhibited photosynthesis (Haydon, 2013). If Mg affects the circadian period when photosynthesis is inhibited by DCMU, and sugars are buffered by 1 % sucrose in the medium, then the effect must occur through pathways not related to photosynthesis.

In the presence of sucrose, DCMU has little or no effect on circadian period (Fig. 5) as reported previously (Haydon et al., 2013; Takahashi et al., 2015). Magnesium deficiency profoundly affected the circadian period in the presence or absence of DCMU when sucrose was added to the medium. Hence, the effect of Mg on the oscillator might not be due to the inhibition of photosynthesis and associated downstream processes (Fig. 5*B,E*). The attempt to examine the effect of DCMU combined with Mg deficiency in the absence of added sucrose resulted in a strong effect of DCMU on plant performance and health making it impossible to detect circadian rhythms (Fig. S5).

**Fig. 5.**
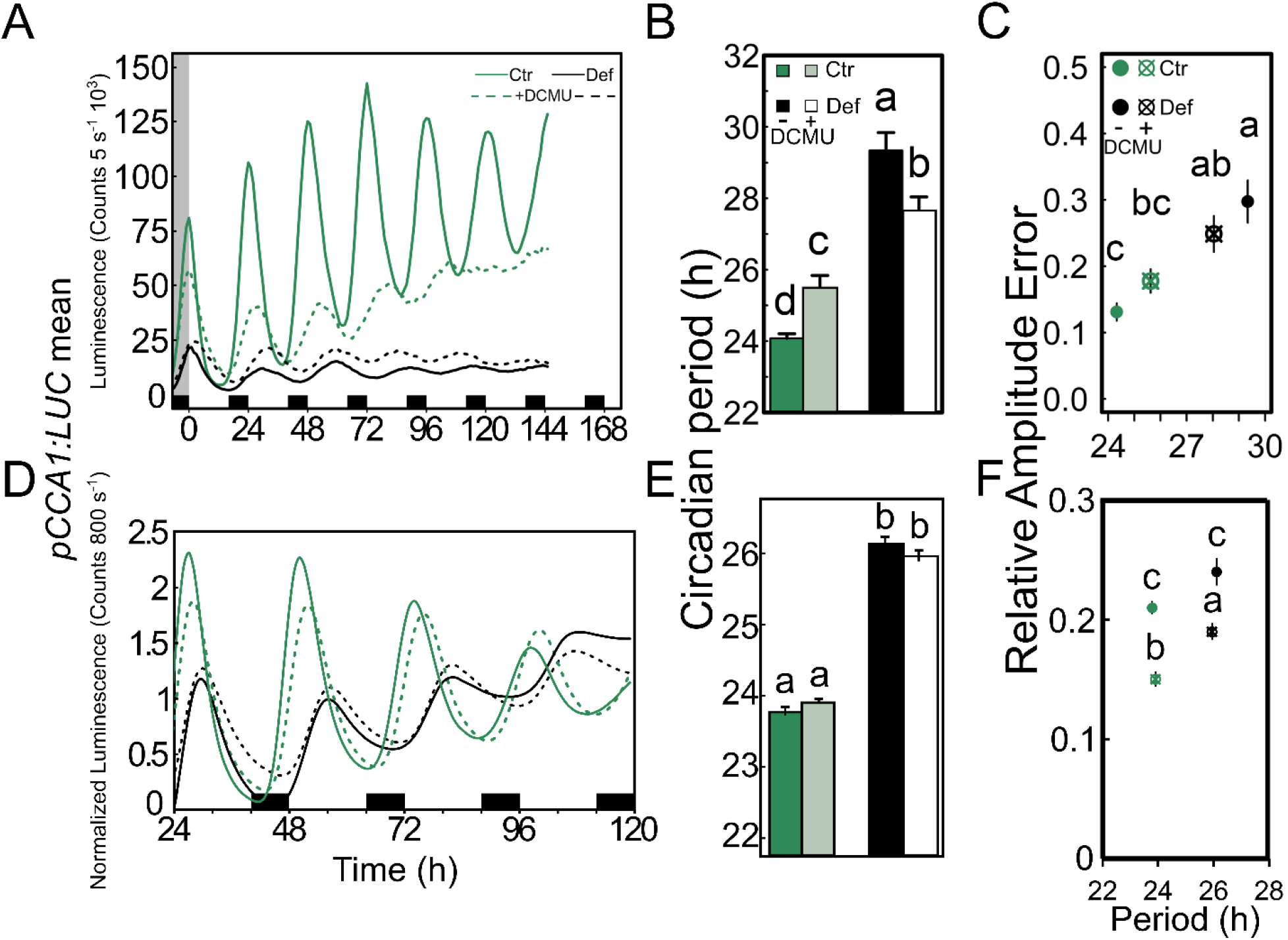
An active photosynthetic system is not required to detect Mg deficiency-dependent circadian alterations. Seedlings entrained for eight days to 16/8-h light/dark cycles on either 200 μM (Ctr) or 5 μM (Def) Mg media were released into LL in the presence or absence of 20 μM DCMU. The experiment was independently repeated [Experiment 1 A-C; Experiment 2 D-F]. (*A,D*) Luminescence traces of *pCCA1:LUC+* reporter in LL, (*B,E*) estimated circadian period (h) of *pCCA1:LUC* activity (mean ± SEM, *n* = 12), (*C,F*) relative amplitude error of *pCCA1:LUC* oscillations (mean ± SEM, *n* = 12). Statistical significance was verified by factorial ANOVA followed by Tukey HSD post-hoc (different letters indicate significance at the level of *P* < 0.05). Experiments were undertaken in the laboratories at Université libre de Bruxelles (*A-C*) and Cambridge University (*D-F*).

### External Mg supply is unlikely to be an entrainment stimulus

To determine whether Mg could act as a clock regulator in plants, a 4 h long pulse was applied with 10 mM Mg in intervals of 3 h to seedlings that had been entrained with 50 μM Mg (insufficient concentration, see Fig. 1) for eight days. Under those conditions, a Mg pulse did not cause a phase shift of *pCCA1:LUC* peak expression at any time point (Fig. 6*A*). It can be concluded that Mg is unlikely to be a zeitgeber for the circadian clock in Arabidopsis. However, the applied Mg pulse decreased the RAE at all the time points (Fig. 6*B*) and full resupply of Mg to deficient seedlings restored rhythmicity independent at which time of day Mg was resupplied (Fig. 6C). When seedlings were grown on medium overly supplied with Mg, there was no effect on circadian period in the presence or absence of 1 % sucrose (Fig. S6). Thus, sufficient Mg supply seems necessary to maintain proper functioning of the circadian oscillator but is not associated with entrainment.

**Fig. 6.**
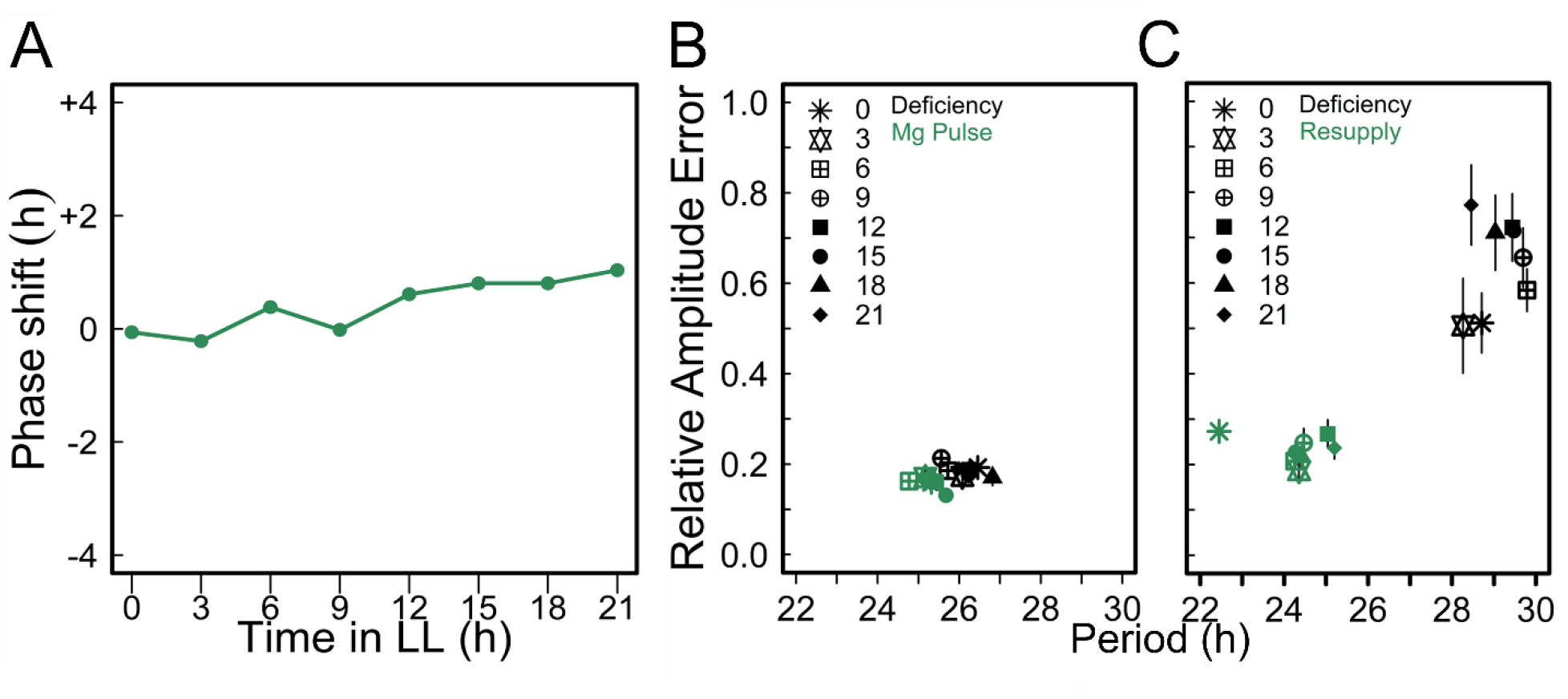
Mg is not a zeitgeber to set circadian time. Seedlings were entrained for eight days to 12/12-h light/dark cycles on medium supplied with 50 μM Mg before released into LL. A pulse of 10 mM Mg was applied during 4 h in 3 h intervals from ZT0 in LL along one circadian cycle. (*A*) Phase response of *pCCA1:LUC* activity rhythms to pulses of Mg at different time points in LL. (*B*) RAE of the oscillations (mean ± SEM, *n* = 6). (*C*) RAE of *pCCA1:LUC* oscillations of seedlings entrained on medium containing 5 μM Mg that were resupplied with 3 mM Mg every 3 hours under LL conditions (means ± SEM, n = 6).

### Simulations in a model of the Arabidopsis circadian oscillator suggests that Mg globally affects the kinetics of the circadian oscillator

A mathematical model (De Caluwé et al., 2016) was used to gain further insight concerning the mechanism underlying the impact of Mg nutrition on the circadian oscillator. With the default parameter values (kinetics rates), the model simulates the behaviour of wild type plants in control conditions assuming sufficient supply with Mg. First, single parameters were changed to see whether the predicted circadian period was comparable to what was observed under Mg deficiency in free-running conditions. When modelling reduced PRR5/TOC1 protein degradation (Fig. S7*A*) or reduced *PRR7/PRR9* RNA synthesis (Fig. S7*B*), higher expression of *CCA1/LHY* was predicted by the model while experimentally, Mg deficiency decreased *CCA1* transcript levels (Fig. 3*C*). Reduced RNA synthesis of the *ELF4/LUX* evening complex lowered the amplitude of *CCA1/LHY* mRNA oscillations but did not predict an increase in circadian period (Fig. S7*C*). These results, together with other simulations of single parameter changes (not shown), did not reproduce simultaneously the decreased amplitude of *CCA1* and the increased period observed experimentally. We then tested if alterations in overall kinetic parameters such as rates of transcription, translation and protein degradation could lead to the observed tendencies. The model predicted very long free-running periods and damped oscillations in response to reduced global rates of mRNA and protein synthesis as well as protein degradation rates (Fig 7*A*) resembling the experimental observations under Mg deficiency (Fig 1 *A*). The model predicted a restoration of amplitude and a decrease in circadian period of *CCA1/LHY* activity rhythms when Mg was reintroduced (Fig 7*A*), which is in accordance with experimental data (Fig. 6*C*) and confirms the necessity of Mg to maintain circadian period in Arabidopsis. Thus, alterations in global rates of transcription, translation and protein degradation are likely to be affected by Mg deficiency, provoking an increase in circadian period.

**Fig. 7.**
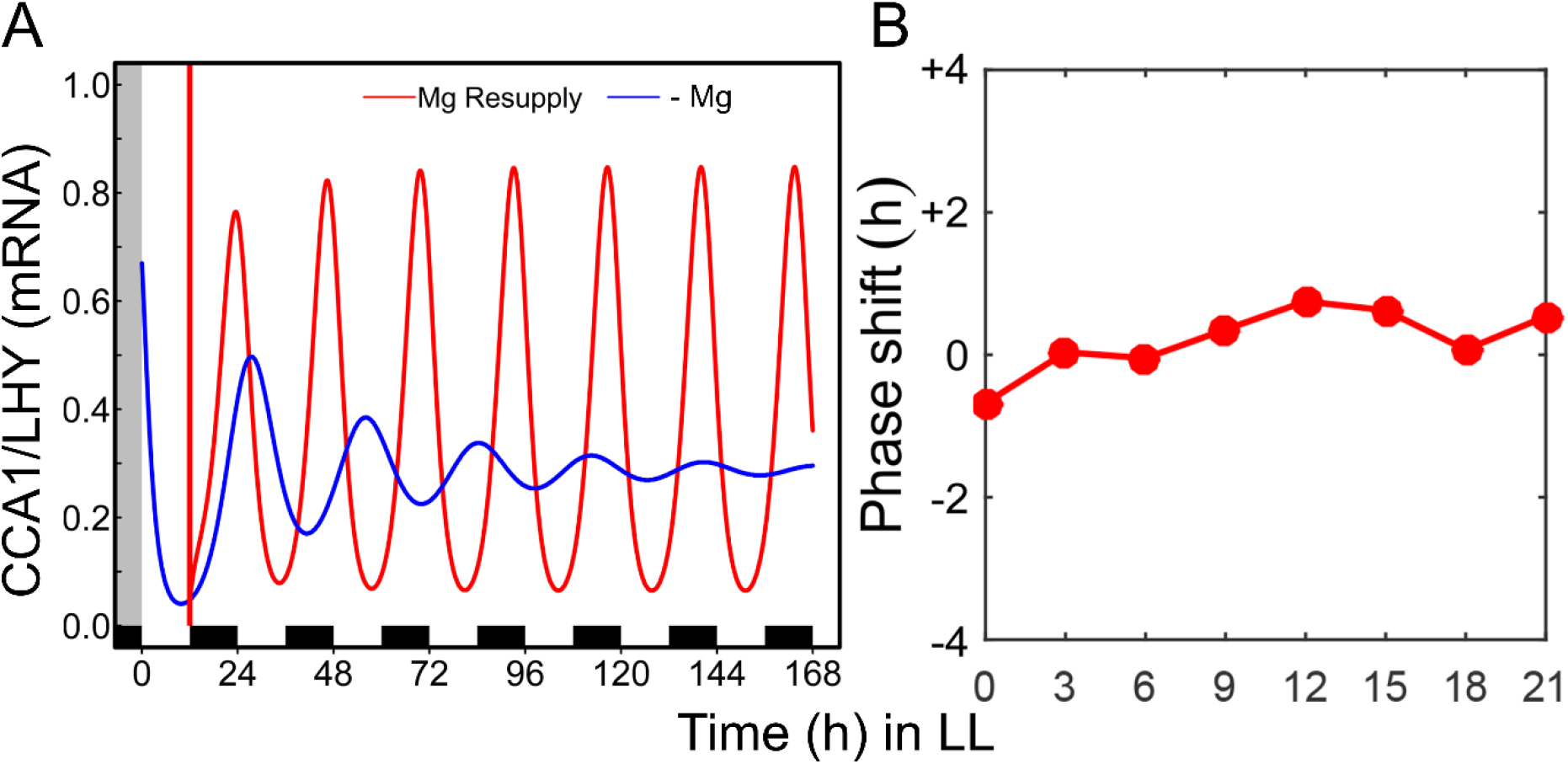
Simulation of global effect of Mg deficiency on transcription/translation rate increases the circadian period of *CCA1* activity rhythms. (*A*) CCA1/LHY oscillations under Mg deficiency (blue) and when Mg was resupplied (red) under the assumption that Mg deficiency affects global rates of transcription and translation. Vertical red bar represents the time of simulated Mg resupply. (*B*) response curve to Mg pulse.

The simulated PRC based on global changes in transcription and translation rate does not predict a significant phase shift in response to a Mg pulse (Fig. 7*B*), which rules out Mg as a zeitgeber for the circadian oscillator and is in line with experimental data (Fig. 6*A*).

A translation inhibition assay using cycloheximide (CHX) was performed combined with different Mg concentrations to test the model prediction that a global decrease in translation could account for the experimental results. The application of 0.5 μg mL^−1^ CHX increased the circadian period of *pCCA1:LUC* and its respective RAE (Fig. 8). The translation inhibitor had a stronger effect on period at low Mg concentrations, independent whether external sucrose was supplied or not (Fig. 8*A,C*). Two-Way ANOVA at a confidence interval of 95 % confirmed a significant correlation between period increased under Mg deficiency and its further increase through translation inhibition when 1 % sucrose was present in the medium (p < 0.05). The synergistic interaction between Mg deficiency and CHX suggest that both affect the circadian system by interfering with translational processes supporting the prediction that Mg deficiency has global effects on kinetics of the circadian oscillator.

**Fig. 8.**
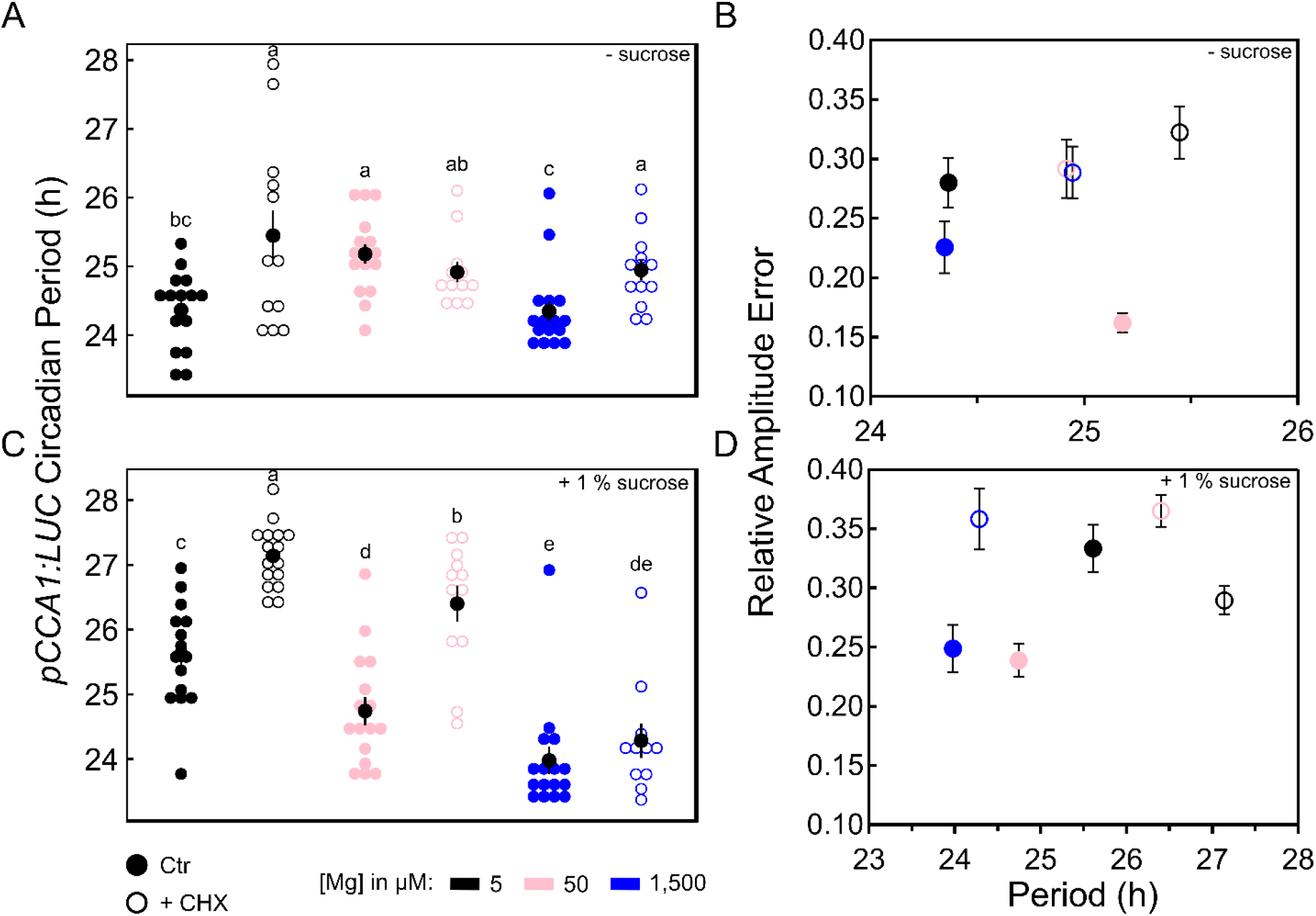
Inhibiting translation increases circadian period of *pCCA1:LUC*. *pCCA1:LUC* Col-0 seedlings were entrained for 11 days to 12 hour light/dark cycle on media supplied with different Mg concentrations before released into LL. Media were either without sucrose (*A-B*) or supplemented with 1 % sucrose (*C-D*). 0.5 μg mL^−1^ CHX (+ CHX) or 0.1 % DMSO as control (Ctr) were applied after 24 hours in LL. Estimated circadian period in hours (h) (*A, C*) and respective RAE (*B, D*) of *pCCA1:LUC* activity under LL after treatment. Significance was determined by a Wilcoxon Rang Sum test. Different letters indicate significance at the level of *P* ≤ 0.05.

## Discussion

Magnesium is essential for multiple processes in plants. It is highly important for photosynthesis where it is bound in the chloroplast as a key compound of the energy transfer in chlorophyll (Lilley et al., 1974; Strasser and Butler, 1977; Walker and Weinstein, 1991). Additionally, it is crucial for sucrose loading into the phloem and its partitioning from source leaves to sink plant organs. Also, Mg is vital to the cellular energy metabolism and sustaining the ribosome structure and is therefore important for protein translation (Chen et al., 2017). In the present study, it was shown that Mg deficiency increased the circadian period of *pCCA1:LUC, pPRR7:LUC* and *pTOC1:LUC* in a dose-dependent manner in Arabidopsis seedlings under constant light and the effect on period was greater when 1 % exogenous sucrose was supplied. (Fig. 1*A*, Fig. S1). Period increase due to Mg deficiency was even more pronounced when seedlings were entrained to long day conditions before release into continuous light (Fig. 4*B*). Within the experimental set-up Mg was supplied as MgSO_4_. Under deficient conditions Na_2_SO_4_ or K_2_SO_4_ were added to the medium to avoid sulphur deficiency following MgSO_4_ restriction. To exclude higher Na^+^ or K^+^ concentrations as being the cause for the observed period increase, MgCl_2_ was used as Mg^2+^ source, which resulted in an increase of circadian period (Fig. S8) confirming Mg depletion being the cause of period increase.

An increase of circadian period caused by iron deficiency was shown to result from disordered photosynthetic functioning (Chen et al., 2013; Salomé et al., 2013). Sugars deriving from photosynthesis feed into the circadian clock, defining a metabolic dawn and adjusting the phase of *CCA1* expression (Haydon et al., 2013). Inhibition of photosynthesis lengthens the circadian period as does constant dim light (Haydon et al., 2013). Here, period lengthening caused by Mg deficiency was dependent on light signalling (Fig. 4) but the observed effects were not due to direct effects of Mg deficiency on photosynthesis. In the presence of DCMU to inhibit photosynthesis and 1 % sucrose to buffer changes in associated sugar production, we found that Mg deficiency had a profound effect on the circadian period of *pCCA1:LUC* oscillations (Fig. 5). Because Mg deficiency could affect the oscillator when photosynthesis was impaired and when changes in the sugar production are buffered, we conclude that Mg might affect the oscillator through other pathways. That the effect of Mg is greater in the presence of added sucrose suggests an association with some energy dependent mechanism.

The phase of the circadian oscillator entrains through signals that regulate its individual components based on their temporal availability. Magnesium deficiency induced a phase delay of *pPRR7:LUC* peak expression in 12/12 h light/ dark cycles (Fig. 2, S3). Interpretation of the timing of LUC signals must be treated with caution as peak of expression is a product of both, promoter activity as well as the rate of LUC translation and folding. However, because the entrained phase of *pPRR7:LUC* was more sensitive to Mg levels than either *pCCA1:LUC* or *pTOC1:LUC* it might be an example of dynamic plasticity of the circadian oscillator in response to its environment (Webb et al., 2019). Dynamic plasticity refers to the plasticity of oscillator period and phase to environmental signals and the ability of the oscillator to keep internal time keeping in synchrony with its environment. Thereby, the individual components of the circadian oscillator are not tightly linked and the relative timing of peak expression between individual components can be plastic up to a certain degree in response to a stimulus (Flis et al., 2016). The effect of *PRR7* timing is notable because while increasing light intensity and different sugars shorten the period in wild-type plants (Farré et al., 2005; Haydon et al., 2013) and nicotinamide lengthens the period (Mombaerts et al., 2019), these responses are absent in *prr7* mutants. The oscillator receives competing signals throughout a cycle of which the “strongest” signal determines the phase of the oscillator. *CCA1* and *TOC1* genes are more locked to dawn and dusk (Fig. 2*A*) making light the determining signal. However, as the oscillator components are not locked to each other, this might assign *PRR7* as the light-independent regulating component in the central oscillator that can respond to external changes and feeds this information into the circadian timekeeping. But Mg does not influence the phase of the oscillator, which excludes Mg as a zeitgeber signal and oscillator phase is strongly determined by light regime. Furthermore, *CCA1* expression is induced by sucrose that sets the phase of *CCA1* peak expression at dawn and is advanced in the presence of exogenous sugars (Haydon et al., 2013). Data in Fig. 2*A* confirm this observation where *CCA1* peaked at an earlier phase in the presence of sucrose under either Mg supply. Sucrose stabilizes circadian oscillations (Haydon et al., 2013; Philippou et al., 2019) as the metabolite enhances the binding of PIF transcription factors to *CCA1* promotor region and correlates with increasing transcript abundance of *CCA1* (Shor et al., 2017).

Circadian oscillations in plants are generated through transcriptional/translational feedback loops whereby sucrose increases translation rates and global protein abundance (Osuna et al., 2007). Proper function of the clock does rely on a diel cycle of transcriptional control (Flis et al., 2016) plus the level of ribosomal loading driven by the circadian clock (Missra et al., 2015). Light induces proteome wide changes in protein abundance in correlation with their transcript abundance depending on the length of photoperiod whereby long days increase the abundance of several photosynthetic proteins that further affected protein abundances of downstream processes (Seaton et al., 2018). Magnesium is a very important co-factor required for translation/ protein synthesis (Chen et al., 2017). It stabilizes ribosomal structure (Klein et al., 2004) and is required for ribosome activity and translation (Weiss and Morris, 1973; Sperrazza and Spremulli, 1983). Inhibiting translation increases the circadian period of *CCA1:LUC* in *Ostreococcus* (Feeney et al., 2016) similar to the effect of Mg depletion reported here under continuous light. Feeney et al. (2016) demonstrated that a high endogenous Mg level increased translation rates in *Ostreococcus* and circadian oscillations of Mg levels in those cells correlate with circadian dependent translation rates. Light-signalling contributes to photoperiod-dependent changes in gene expression at dawn because of the impact of light on transcript abundance (Flis et al., 2016). An interference of transcriptional and translational processes by Mg deficiency might lengthen the period of the circadian oscillator in Arabidopsis in free running conditions where dawn and dusk are absent as strong entrainment signals. This is in accordance with a simulated increase in period obtained by a mathematical model assuming that Mg deficiency impacts overall kinetic parameters like translation rate, transcription rate and rates of protein degradation (Fig 7*A*). In support of the model’s prediction, low doses of the translation inhibitor CHX synergistically increased the circadian period of *pCCA1:LUC* activity with Mg deficiency (Fig. 8), which is in line with the results obtained in *Ostreococcus* and human cells (Feeney et al., 2016). Modelling Mg resupply to deficient seedlings decreased the circadian period as well as restored rhythmicity (Fig 7*A*) and was also shown experimentally (Fig. 6*C*). Under excess Mg supply a period length of nearly 24 h was maintained (Fig. S6). Both underline the importance of Mg as a crucial cofactor for processes related to circadian timekeeping such as transcriptional/translational control.

## Conclusion

Magnesium is essential for proper timekeeping in *Arabidopsis thaliana*. We find that insufficient Mg supply increases circadian period and causes phase delay in light/ dark cycles. Our data suggest that one mechanism by which Mg deficiency can affect the oscillator might be through interference with global translational/ transcriptional processes. Whilst Mg deficiency can affect circadian function we find no evidence that changes in Mg levels can act as zeitgeber setting circadian time.

## Material and Methods

### Plant material and growth conditions

*Arabidopsis thaliana* wild type seeds and reporter lines *pCCA1:LUC, pPRR7:LUC* and *pTOC1:LUC* were all in Columbia-0 ecotype background. Detailed description about luciferase-expressing construct is available elsewhere (Salomé and McClung, 2005). Seeds were sterilized as previously described (De Caluwé et al., 2017), individually plated on self-prepared Murashige and Skoog medium adapted from Hermans et al. 2010a, solidified with 0.5 % Mg-free high gel strength agar (Sigma-Aldrich, Germany) and supplied with different concentrations of Mg (Table S1). Thereby, no sucrose or 1 % sucrose was added to the media, respectively. After stratification for two days in the dark at 4°C, seedlings were entrained for 8-11 days to different photoperiods with white light (~ 100 μmol photons m^2^ s^−1^) under constant temperature of 19°C (Panasonic MLR-352-PE, Netherlands). Seedlings were entrained to the respective Mg concentration from germination on unless stated otherwise. Treatment with DCMU was done as described elsewhere (Haydon and Webb, 2016). For PRC, Mg-deprived seedlings (50 μM) were transferred to medium containing 10 mM Mg for 4 h in 3 h intervals and subsequently returned to entrainment medium. For the Mg resupply assay, Mg deprived seedlings were resupplied in 3 h intervals by transferring seedlings to medium containing 3 mM Mg (Fig. S9). The PRC was calculated as descripted previously (Johnson, 1992).

### Ionome profiling

Arabidopsis Col-0 seedlings were cultivated, as described above, on media supplemented with either 200 μM or 5 μM Mg from germination on. Fifteen pooled seedlings were harvested in the morning (ZT4) on days 8 to 12, rinsed three times in deionized water, carefully cleaned for left-over agar medium and dried at 70 °C for 72 h. Dried plant material was digested in 500 μL 35% (v/v) HNO_3_ at 90 °C for 1 h. From the digest, 200 μL was diluted in 7 mL Milli-Q water and filtered. The mineral concentrations were determined by NexION 350S ICP-MS (PerkinElmer). Indium was used as internal standard to correct for instrument instability and oxide interferences.

### Luciferase experiments and rhythms analysis

Seedlings were sprayed with 2 mM Luciferin (VivoGlo™ Luciferin, In Vivo Grade, Promega, Netherlands) one day before they were released either into continuous light or dark. Following the dosing, seedlings were individually transferred to 96-well opaque white microplates (23300, Berthold, Germany) containing 150 μL of the respective liquid growth medium plus 2 mM D-Luciferin. Microplates were sealed with transparent EASYseal sealing film 120×80 mm (Greiner bio-one, Germany) that was punctured (~1.0 mm Ø) to allow gas exchange. Luminescence was detected at 590 nm for 5-10 s hourly after 120 s in darkness using a multimode microplate reader (TriSta^r2^ LB942 Berthold, Germany). Data were processed with MikroWin^TM^ software v. 5.21 (Labsis Laborsysteme, Germany) and rhythms of LUC activity were analysed with BioDare2 beta (https://biodare2.ed.ac.uk/) (Zielinski et al., 2014). Period and RAE estimates were calculated on rhythms between 24-120 h in continuous light on non-normalized data using Fast-527 Fourier Transformed Non-Linear Least Squares (FFT-NLLS) after linear detrending.

### RNA Isolation and RT-qPCR

Total RNA was isolated from 100 mg frozen ground tissues of whole seedlings. RNA was purified with Maxwell^®^ 16 LEV Plant RNA Kit (Promega, Benelux BV) using Maxwell^®^ 16 AS2000 Instrument (Promega, Benelux BV) according to the manufacturer’s recommendation. Quality and purity of the samples were verified with a NanoDrop 2000 UV-Vis Spectrophotometer (Thermo Scientific, Loughborough, UK). cDNA was synthesized from one μg RNA with GoScript™ Reverse Transcription System (Promega, Benelux BV) and thereafter diluted to 1:30 (v/v) with autoclaved nuclease-free water for quantitative real time PCR (qPCR). qPCR was carried out in 96-well microplates in the PikoReal real time PCR system (Thermo Scientific, Loughborough, UK). Each reaction contained 5 μL of 2x SYBR Green mastermix (Promega, Benelux BV), 2.5 μL primer mix (forward and reverse, 2.5 μM each) and 2.5 μL 1:30 diluted cDNA. Thermocycles were as followed: pre-incubation at 95°C for 3 min, 40 cycles at 95°C for 30 s, 60°C for 1 min. *ELONGATION FACTOR 1α (EF1α*, At5g60390), *UBIQUITIN 10 (UBQ10*, At4g05320) and *CYCLIN-DEPENDENT KINASE A;1 (CDKA*, At3g48750) were used as reference genes. Oligo nucleotide sequences and their respective efficiencies are given in Table S2.

### Mathematical Modelling

A previously developed model (De Caluwé et al., 2016) of the core oscillator was used to investigate the mechanisms of Mg input to the clock. The model consists of eight differential equations describing the evolution of mRNA and protein concentrations of four pairs of clock genes: CCA1/LHY, PRR9/PRR7, PRR5/TOC1, and ELF4/LUX. The kinetic parameters represent synthesis rates, degradation rates, and enzymatic constants. The full equations and parameter values for the model are described elsewhere in detail, along with a description of the building and optimization process (De Caluwé et al., 2016). We used the original parameter values to represent plants fully supplied with Mg. The intermediate- and low-Mg conditions were modelled by simulating by reducing the mRNA and protein synthesis rates by a factor of 0.5 (intermediate-Mg) or 0.35 (low-Mg), and the protein degradation rates by 0.7 (intermediate-Mg) or 0.65 (low-Mg). The periods of the Mg-deficient clock were around 26 h (intermediate-Mg) or 28.5 h (low-Mg). For modelling single parameters as an Mg input, we multiply specific kinetic parameters by 0.2 (representing a reduction of 80% of their value).

To construct the PRC, entrainment cycles were simulated on either intermediate-Mg or low-Mg conditions and, thereafter, Mg resupply was simulated in continuous light by restoring all parameters to their initial values. The circadian phase was calculated as previously described (Johnson 1992). Phase shifts were defined as the difference in circadian phase between the first peaks of the treated and control simulations. All simulations were performed in MATLAB (Mathworks, Cambridge). The integration of the differential equations used the external CVODES solver.

### Translation inhibition assay

For the assay, clusters of 8-10 *pCCA1:LUC* Col-0 seeds were sown in plastic rings sealed at the base with 0.5 μm nylon mesh on solid media (0.5 % Mg-free agar) containing different concentrations of Mg either in the presence or absence of 1 % sucrose. Growth conditions were the same as described earlier. Eleven days old seedlings were dosed with 2 mM Luciferin and luminescence was detected for 800 s hourly under LL with a Nightshade CCD camera and imaging chamber (Berthold). The translation inhibitor was applied just before subjective dawn of the second day LL by transferring seedlings to liquid media with the respective Mg supply containing 0.5 μg mL^−1^ CHX (dissolved in DMSO) or 0.1 % DMSO as solvent control. Captured data were processed with IndiGO software (Berthold) and rhythms of *pCCA1:LUC* were analysed with BioDare2 beta (https://biodare2.ed.ac.uk/) as descripted earlier. Replicates with a RAE ≥ 0.5 were excluded from statistical analysis. A two-way ANOVA was performed followed by Wilcoxon Rang Sum test to determine whether CHX and/ or Mg deficiency significantly alter the circadian period with a confidence interval of 95 %.

### Statistical analysis

All statistical analyses performed in this study were carried out in R software version 3.4.1. First, the distribution of the residuals was checked using the functions *hist* and *shapiro.test* (packages *graphics* and *stats*, respectively). Homogeneity of variances across groups was tested with Levene’s test (*leveneTest* from package *car*). Parametric statistics were performed (in experiments with normally distributed and homoscedastic residuals) by using Two-Sample Student’s *t*-test with 95% confidence interval (*t.test* function from package *stats*) to compare the means between two groups of values. One-way ANOVA or factorial ANOVA were used to compare the means between more than two groups (*aov* function from package *stats*). Tukey’s ‘Honest Significant Difference’ (HSD) method was performed to generate a confidence interval (C.I.) on the differences between multiple means being compared with either ANOVA tests (*TukeyHSD* function from package *stats*). For non-parametric statistics, Kruskal-Wallis Rank Sum test followed by Nemenyi post-hoc test were performed (*kruskal.test* and *posthoc.kruskal.nemenyi.test* functions from packages *stats* and *‘PMCMR’*, respectively).

## Acknowledgments

We thank Dr. C. Robertson McClung (Dartmouth College) for providing the luciferase reporter seeds. We are in debt with Dr. Natsuko Kobayashi and Keitaro Tanoi (University of Tokyo) for the ICP-MS facility. We thank Dr. Michael Haydon (University of Melbourne) and Dr. John O’Neill, for insightful discussions about luminescence and PRC data.

## SI Tables and Figures

**Table S1.**
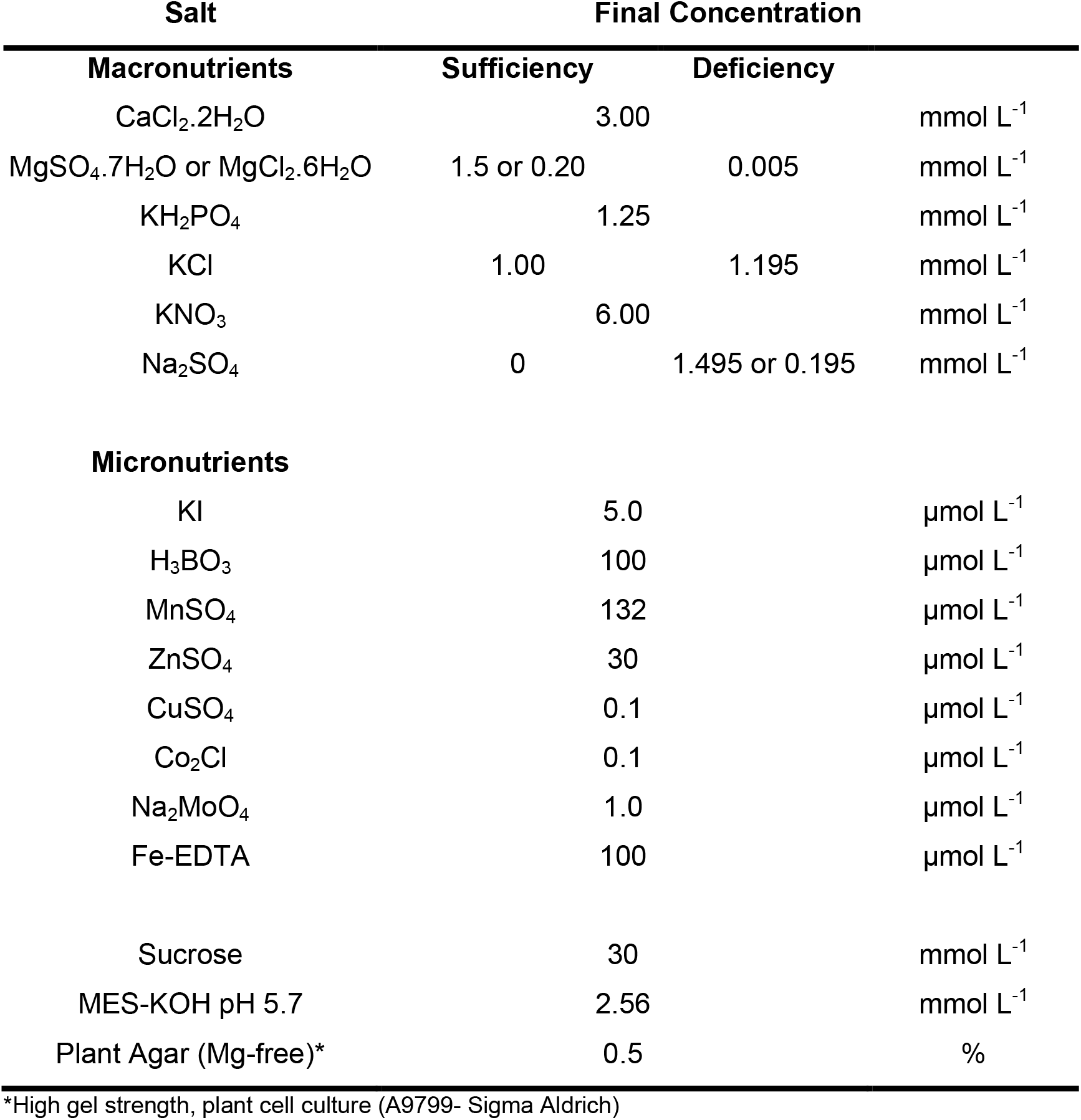
Composition of modified MS medium for *in vitro* plant culture.

**Table S2.**
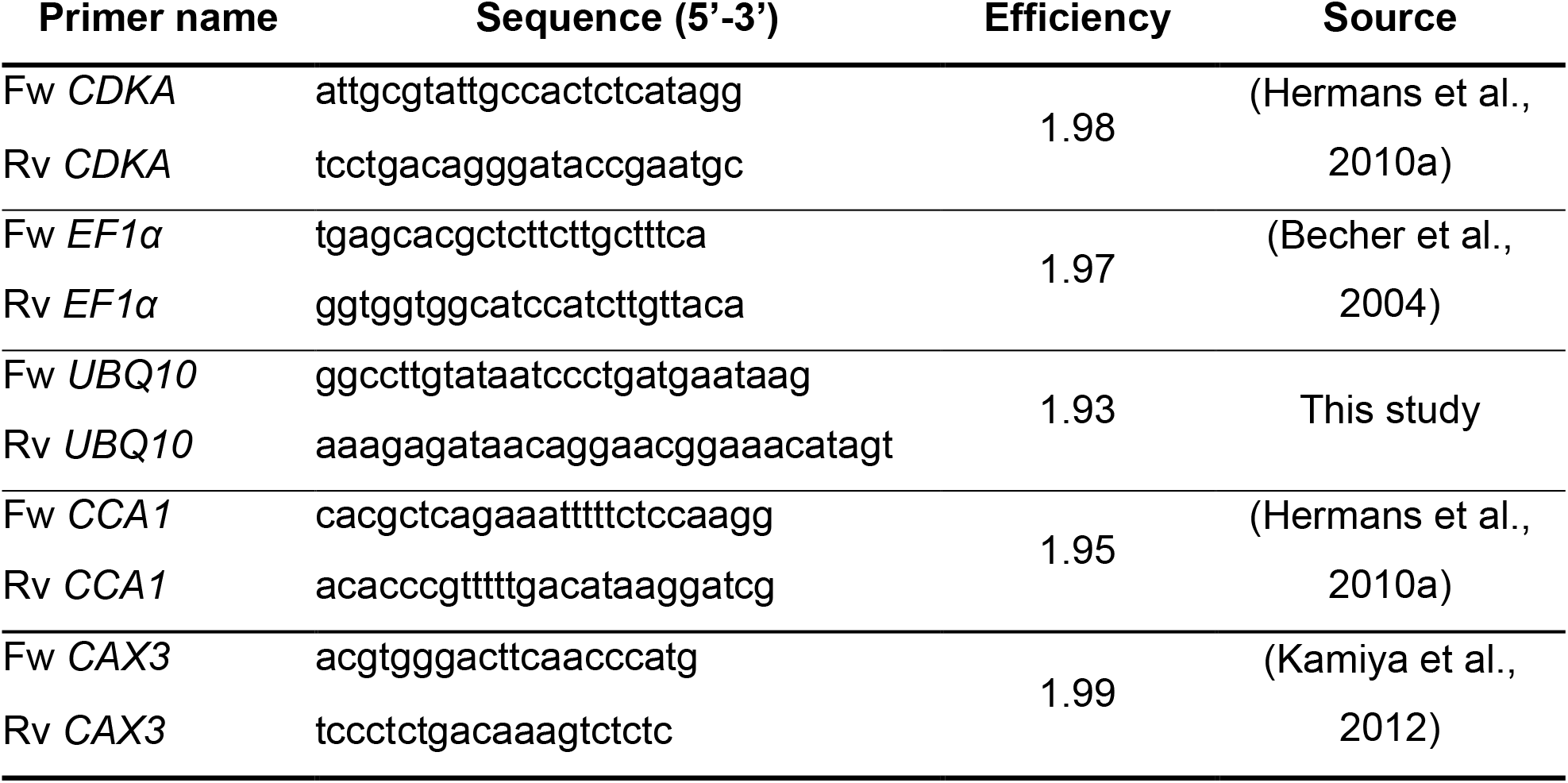
List of primers used for quantification of transcript levels.

**Fig. S1.**
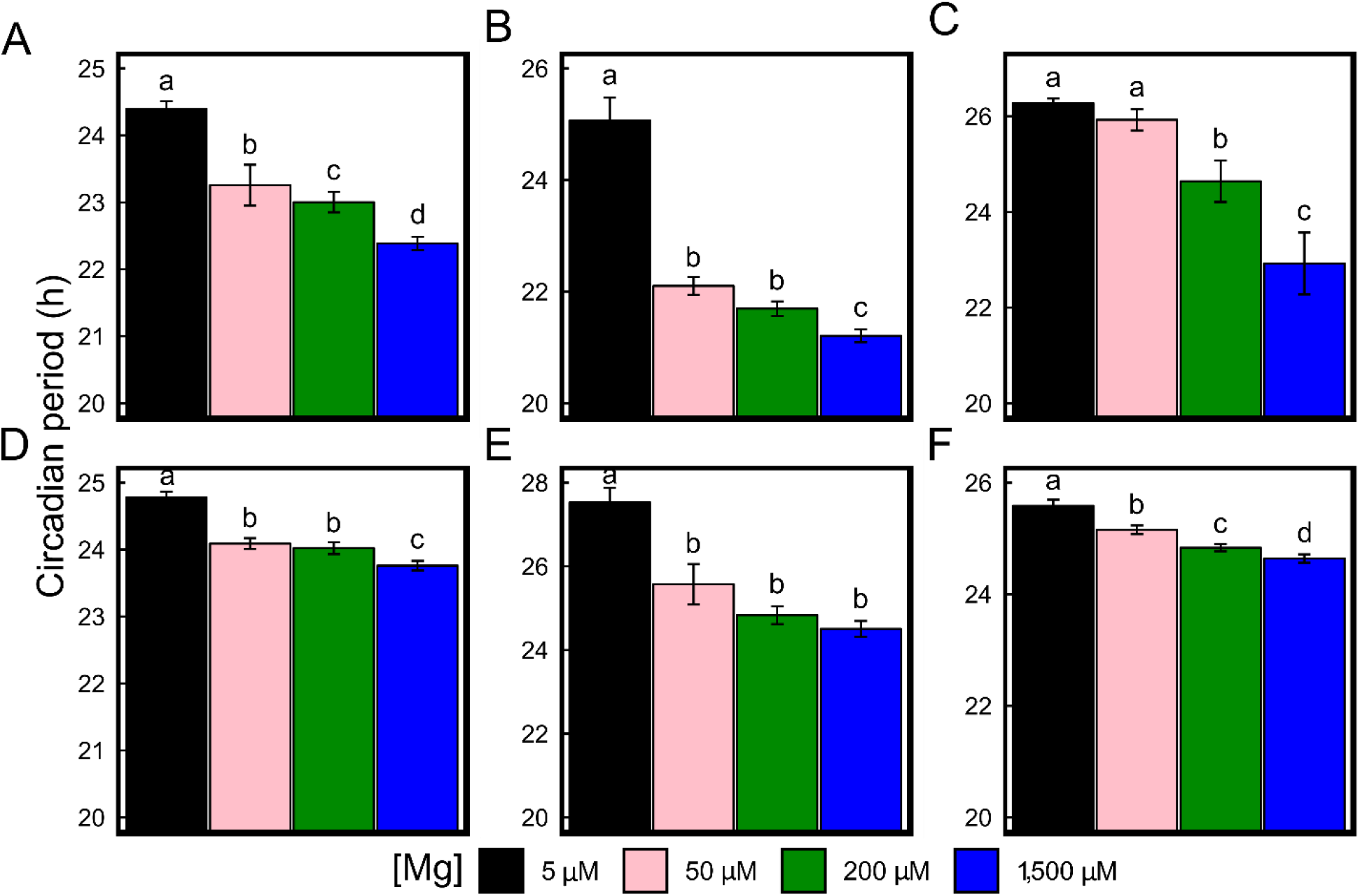
Limiting Mg availability increases the circadian period in *Arabidopsis thaliana* seedlings. Estimated circadian period in hours (h) under continuous light from different Col-0 LUC report lines of the central oscillator (mean ± SEM, n = 20). Seedlings were entrained for 11 days to 12 hour light/dark cycle on media supplied with different Mg concentrations and released into continuous light. Media were either supplemented with 1 % sucrose (A-C) or without sucrose (D-F). (A, D) *pCCA1:LUC*, (B, E) *pPRR7:LUC*, (C, F) *pTOC1:LUC*. Significance was determined by a Wilcoxon Rang Sum test. Different letters indicate significance at the level of *P* ≤ 0.05. Experiments were undertaken indepently in laboratories of Cambridge University in oder to repeat results obtained at Université libre de Bruxelles (Fig. 1) and to complement those data using different reporter genes of the circadian oscillator.

**Fig. S2.**
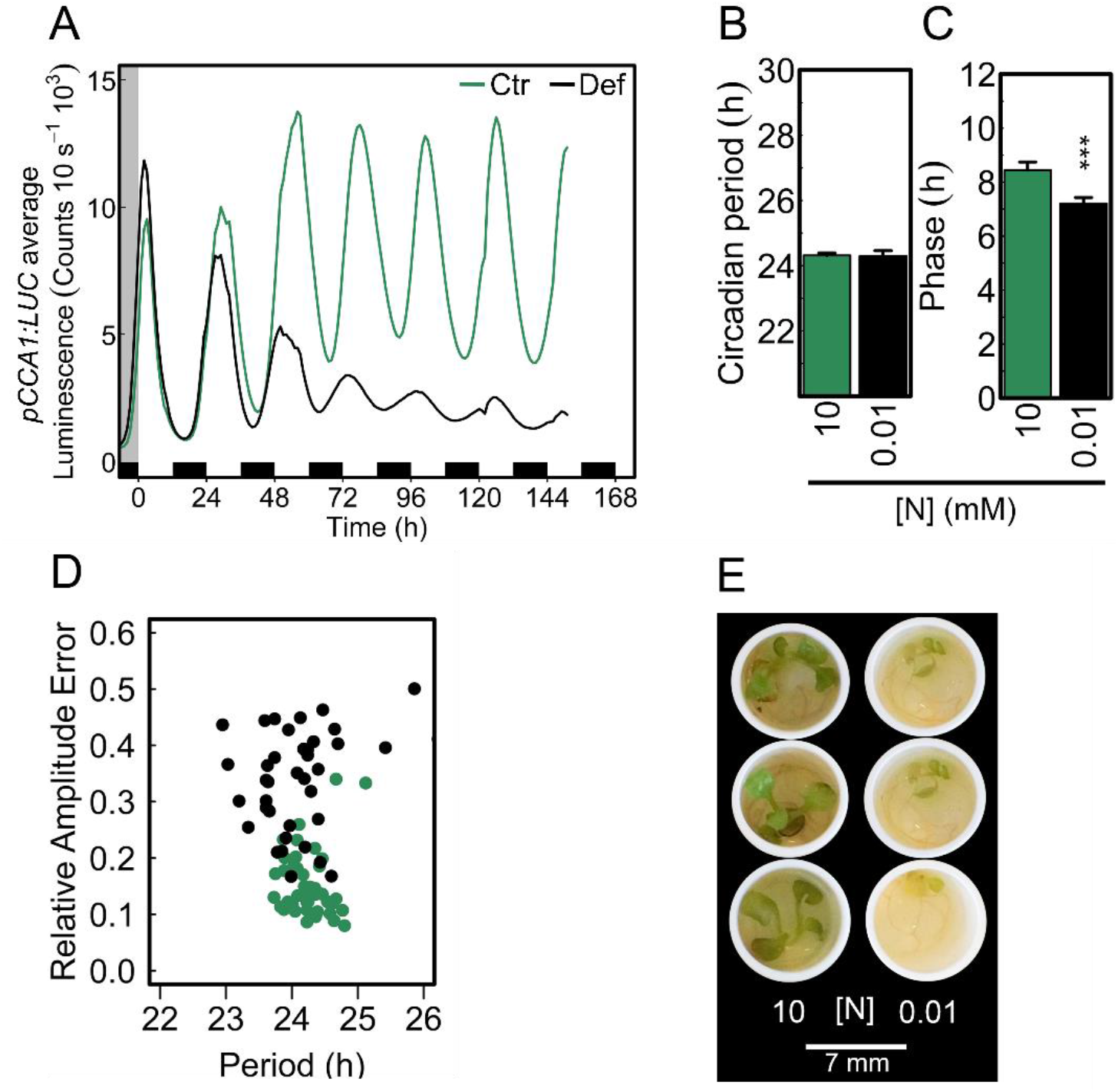
Limited external N availability does not alter the circadian period. Seedlings entrained for eight days to 12/12-h light/dark cycles on either control (N [10 mM] supplied as KNO_3_) or N-deficient (N [0.01 mM]; K^+^ shortage was compensated by 9.99 mM KCl) media were released into continuous light. (*A*) luminescence traces of *pCCA1:LUC* activity (*B*) estimated circadian period of *pCCA1:LUC* oscillations under LL, (*C*) circadian phase of *pCCA1:LUC* peak expression, (*D*) relative amplitude error of the oscillations (mean ± SEM, *n* = 48), (*E*) morphological phenotype of seedlings after 7 days in continuous light. Statistically significance was verified by Two-Sample Student’s *t*-test with a 95% confidence interval. Asterisks represent significance at *** *P* < 0.001).

**Fig. S3.**
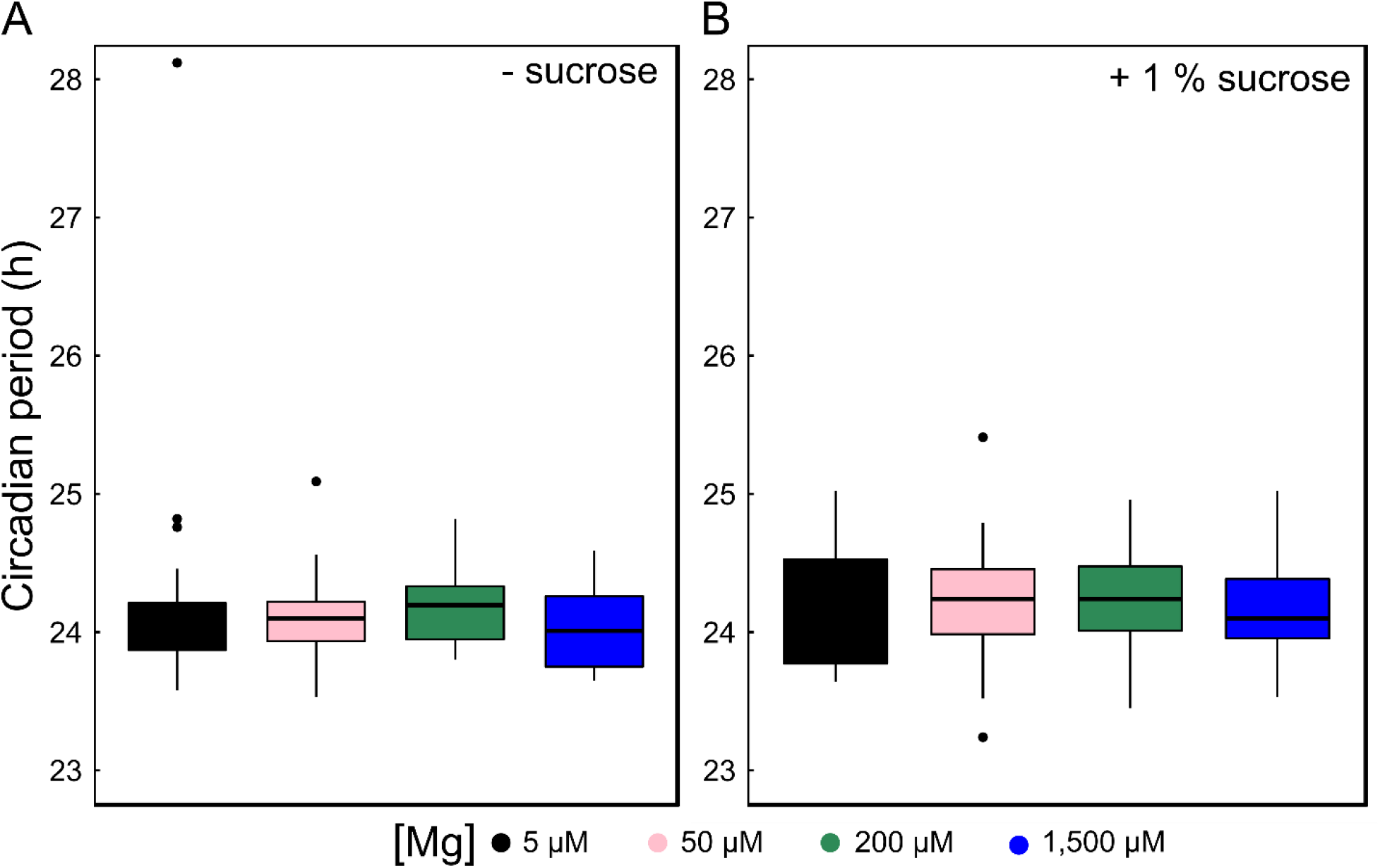
Mg deficiency does not alter the circadian period of *PRR7:LUC* activity under light/dark conditions. Arabidopsis seedlings were entrained for 11 days to 12/12 hour light/dark cycle on media supplied with different Mg concentrations and circadian period of *pPRR7:LUC* activity was estimated over a course of four days under entrainment conditions. (*A*) absence of sucrose, (*B*) presence of 1 % sucrose.

**Fig. S4.**
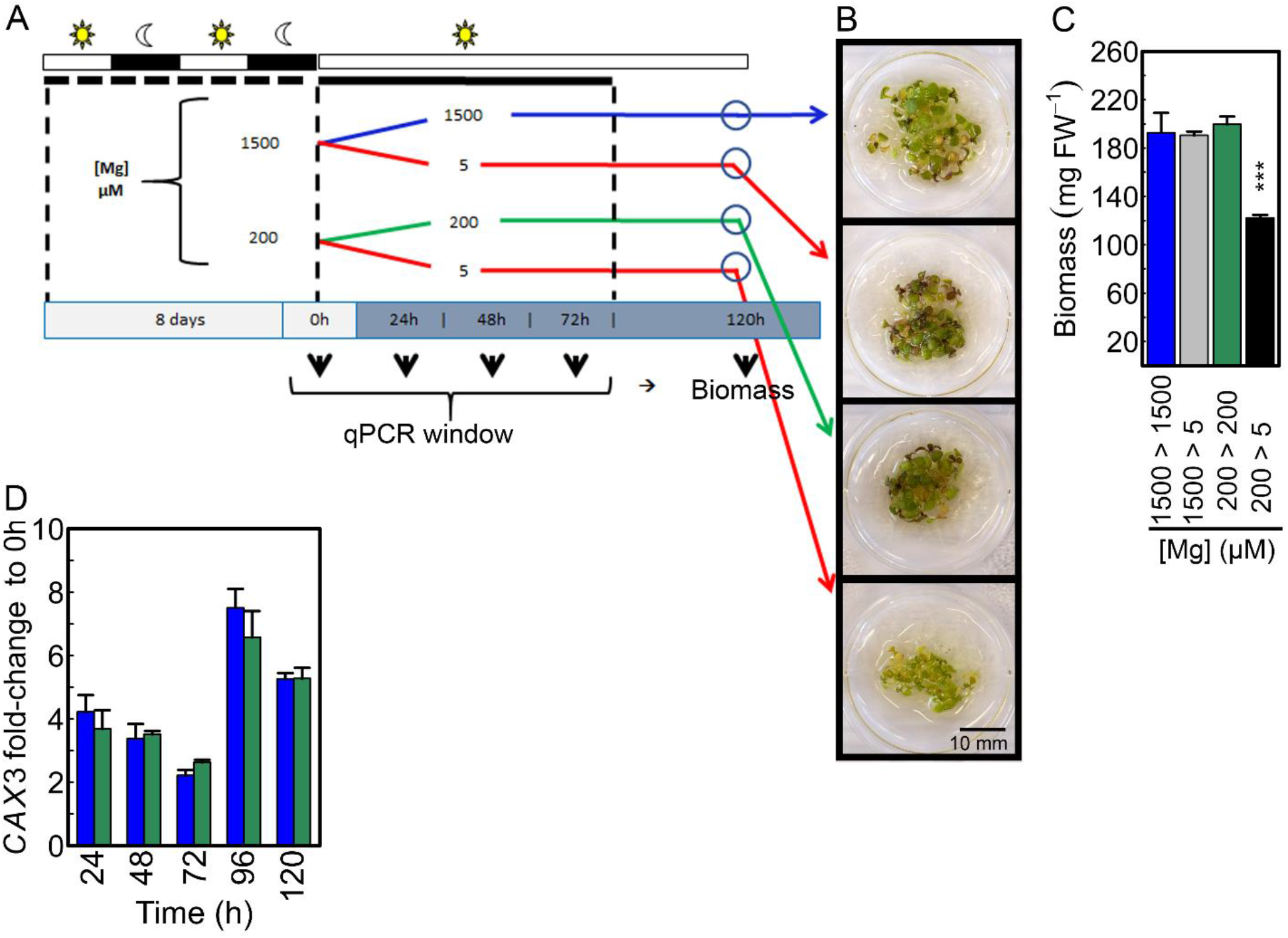
Young Arabidopsis seedlings successfully cope with lower Mg concentrations than the usually supplied in the growth medium. (*A*) Experimental design for seedlings entrained for eight days to 12/12-h light/dark cycles on either 1,500 μM or 200 μM Mg media and thereafter released into continuous light with either fully supplied Mg or Mg-deficient media to assess (*B*) morphological phenotype, (*C*) fresh biomass [mean ± SEM, *n* = 4 (1 = 20 pooled seedlings)], (*D*) mRNA levels of *CAX3*. Statistically significance was verified by One-Way ANOVA followed by Tukey HSD post-hoc test (*P* < 0.001).

**Fig. S5.**
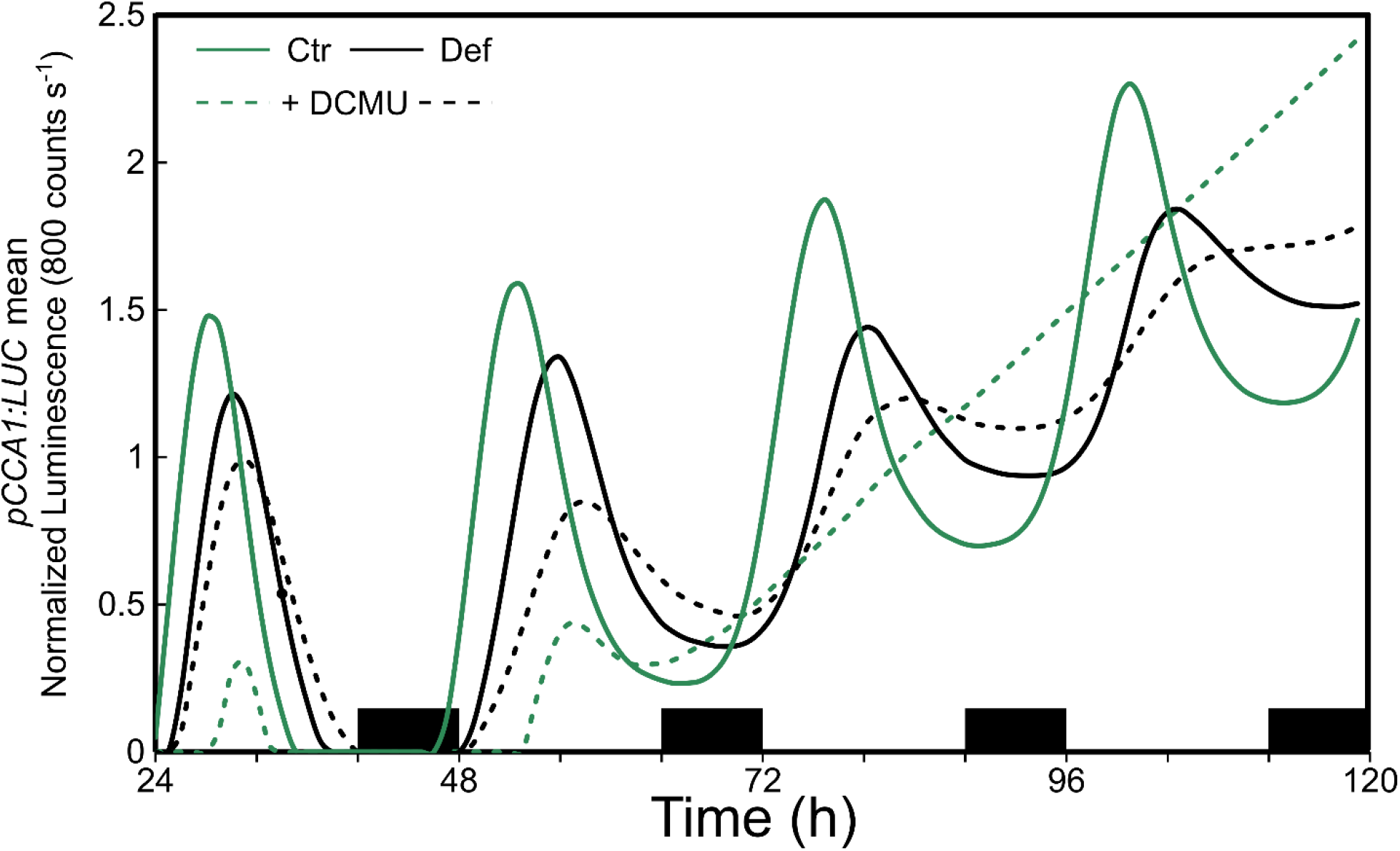
DCMU treatment without external supply of sucrose hampers circadian oscillations of *pCCA1:LUC*. Normalized mean luminescence traces of *pCCA1:LUC* in LL conditions after seedlings were entrained for 11 days to 16/8-h light/dark cycles on either 200 μM (Ctr) or 5 μM Mg (Def) without external supply of sucrose. Seedlings were transferred to respective medium containing 20 μM DCMU prior to release into LL.

**Fig. S6.**
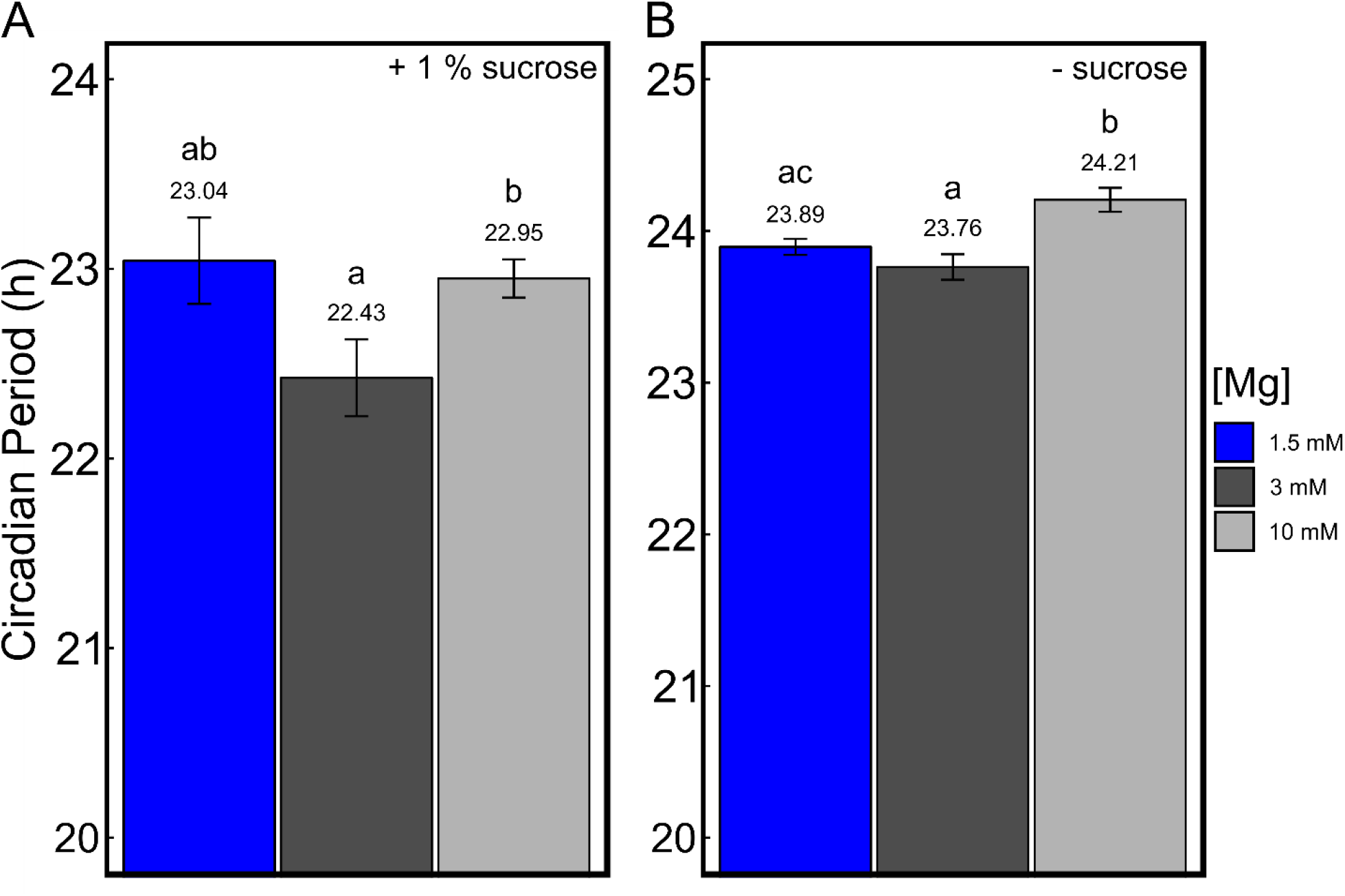
Excess Mg supply maintains the circadian period in *Arabidopsis thaliana* seedlings. Estimated circadian period in hours (h) under LL from *pCCA1:LUC* reporter line (mean ± SEM, n = 20). Seedlings were entrained for 11 days to 12/12 hour light/dark cycle on media supplied with different Mg concentrations before released into LL. Media were either supplemented with 1 % sucrose (*A*) or without sucrose (*B*). Significance was determined by a Wilcoxon Rang Sum test. Different letters indicate significance at the level of *P* ≤ 0.05.

**Fig. S7.**
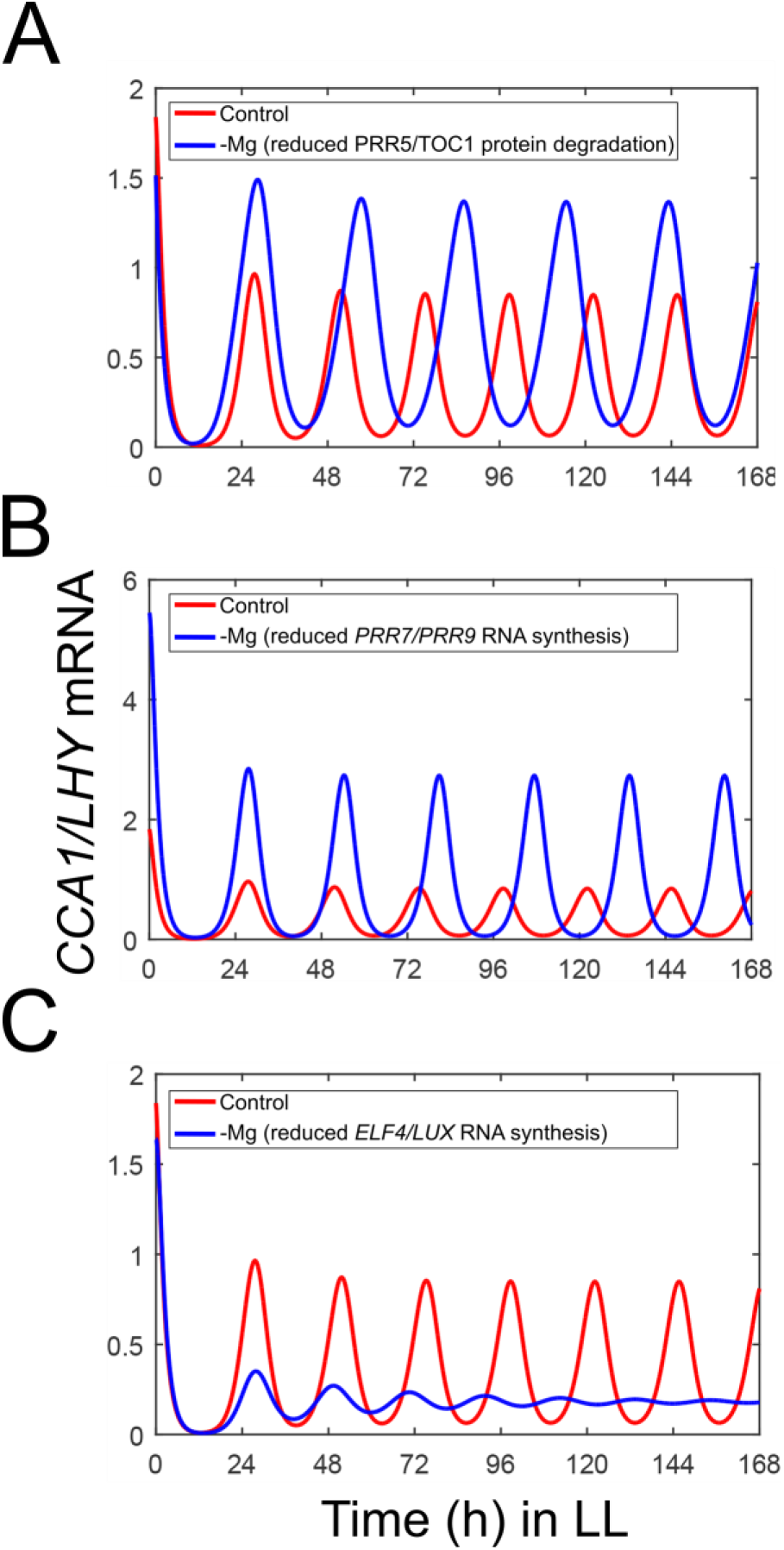
Modelling predictions and translation inhibition assay. Simulations of the behaviour of wild type Arabidopsis circadian clock under free running conditions in LL when single parameters are changed. (*A*) reduced PRR5/TOC1 protein degradation, (*B*) reduced *PRR7/PRR9* RNA synthesis, (*C*) reduced RNA synthesis of the *ELF4/LUX* evening complex.

**Fig. S8.**
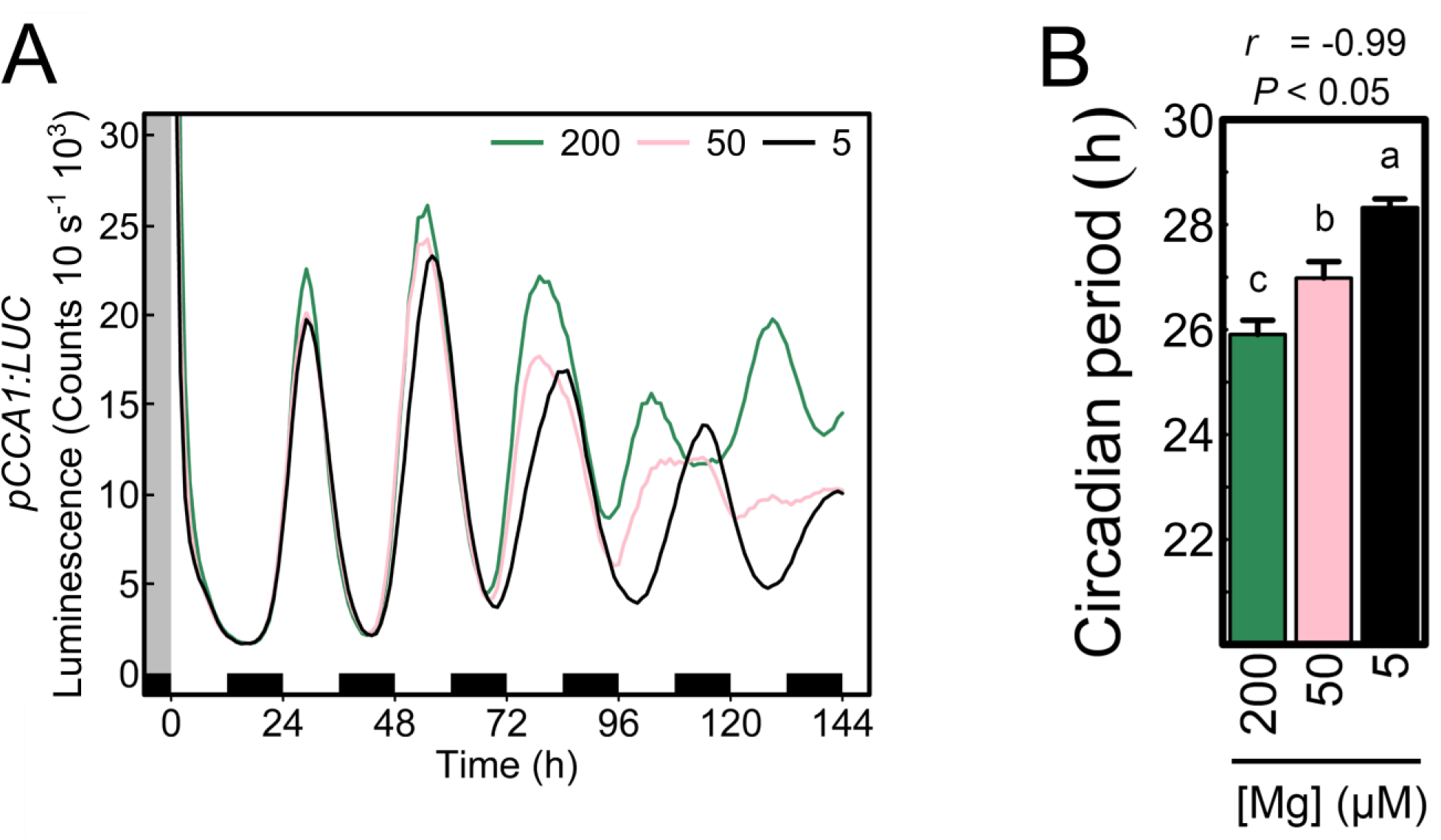
Adding MgCl_2_ as external Mg source increased the circadian period of *pCCA1:LUC*. Arabidopsis seedlings were entrained for eight days to 12/12 hour light/dark cycles on media supplied with the referred Mg concentrations before released into LL. (*A*) average luminescence traces of *pCCA1:LUC* activity, (*B*) estimates circadian period (h) of *pCCA1:LUC* activity (mean ± SEM, *n* = 24) and correlation between circadian period *vs*. external Mg concentrations. Significance was verified by Two-Sample Student’s *t*-test 95% confidence interval, One-Way ANOVA followed by Tukey HSD post-hoc test and Pearson’s rho correlation coefficient. Different letters indicate significance at the level of *P* < 0.05.

**Fig. S9.**
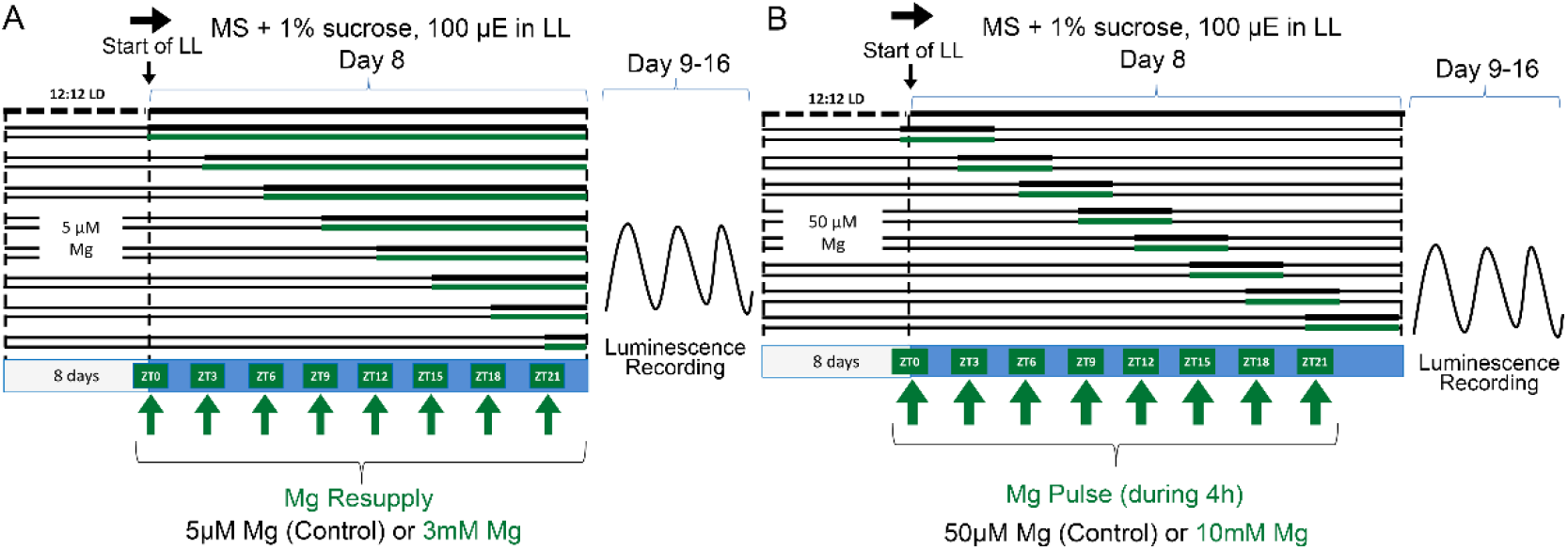
Experimental design of (*A*) Mg resupply and (*B*) phase response curve.

